# The *Histoplasma capsulatum DDR48* Gene Is Required For Survival Within Macrophages, Response To Oxidative Stress, And Resistance to Antifungal Drugs

**DOI:** 10.1101/2020.10.25.354308

**Authors:** Logan T. Blancett, Kauri A. Runge, Gabriella M. Reyes, Lauren A. Kennedy, Sydney C. Jackson, Sarah E. Scheuermann, Mallory B. Harmon, Jamease C. Williams, Glenmore Shearer

**Affiliations:** Department of Cellular and Molecular Biology, The University of Southern Mississippi, Hattiesburg, MS, USA; Mississippi INBRE Research Scholars, The University of Southern Mississippi, Hattiesburg, MS, USA; Mississippi INBRE Research Scholars, Southwest Mississippi Community College, Summit, MS, USA; Mississippi INBRE Research Scholars, Tougaloo College, Jackson, MS, USA

## Abstract

*Histoplasma capsulatum (Hc)* is a systemic, dimorphic fungal pathogen that affects upwards of 500,000 individuals in the United States annually. *Hc* grows as a multicellular mold at environmental temperatures; whereas, upon inhalation into a human or other mammalian host, it transforms into a unicellular, pathogenic yeast. This manuscript is focused on characterizing the DNA damage-responsive gene *HcDDR48*. *HcDDR48* was originally isolated via a subtractive DNA library enriched for transcripts enriched in the mold-phase of *Hc* growth. Upon further analysis we found that *HcDDR48* is not just expressed in the mold morphotype, but both growth programs dependent upon the environment. We found that *HcDDR48* is involved in oxidative stress response, antifungal drug resistance, and survival within resting and activated macrophages. Growth of *ddr48Δ* yeasts was severely decreased when exposed to the reactive oxygen species generator paraquat, as compared to wildtype controls. We also found that *ddr48Δ* yeasts were 2-times more sensitive to the antifungal drugs amphotericin b and ketoconazole. To test *HcDDR48*’s involvement *in vivo*, we infected resting and activated RAW 264.7 murine macrophages with *Hc* yeasts and measured yeast survival 24-hours post-infection. We observed a significant decrease in yeast recovery in the *ddr48Δ* strain compared to wildtype *Hc* levels. Herein, we demonstrate the importance of maintaining a functional copy of *HcDDR48* in order for *Hc* yeasts to sense and respond to numerous environmental and host-associated stressors.

**Importance:** *Histoplasma capsulatum* is an intracellular pathogen of phagocytes, where it subverts immune recognition and avoids killing by the innate immune system. Macrophages provide a permissive environment for *Hc* replication and killing only occurs upon the onset of the T-cell driven adaptive immune response. *Hc* has evolved numerous virulence factors that aid in its survival against host-derived ROS and RNS *in vivo*. While these virulence factors have been described in past years, only a few reports describing the regulation of these genes and how this intricate system leads to fungal survival. In this study, we characterized the stress response gene *DDR48* and determined it to be indispensable for *Hc* survival within macrophages. *HcDDR48* regulates transcript levels of superoxide dismutases and catalases responsible for detoxification of ROS and contributes to antifungal drug resistance. Our studies highlight *DDR48* as a potential target to control *Hc* infection and decrease the severity of the disease process.

## Introduction

*Histoplasma capsulatum (Hc)* is the etiological agent of histoplasmosis, one of the leading endemic mycoses in the world. *Hc* has worldwide distribution, but is primarily endemic to the continents of North America, Central America, and Africa (1, 2). In the United States, *Hc* is found primarily in the MS and OH river valley regions, where it is found in close association with soils enriched with bird or bat guano (3–5). Serological data indicates that roughly 80% of the population within these endemic regions have been exposed to *Hc*, with over 500,000 new cases diagnosed annually (6). *Hc* is a thermally dimorphic fungus, meaning its lifecycle exists in two distinct, temperature-dependent, forms. At environmental temperatures (25°C) the fungus grows as a multicellular, saprophytic mold that produces vegetative microconidia and macroconidia. When soil contaminated with *Histoplasma* conidia is disturbed, the conidia are aerosolized where they are potentially inhaled into a human or other mammalian host’s lungs. The increase in temperature (37°C) within the host’s lungs triggers a transcriptional growth program in *Histoplasma* that promotes a dimorphic shift to unicellular, pathogenic yeasts (3, 7, 8). The dimorphic shift from mold to yeast is critical for *Histoplasma* pathogenesis, as locking *Hc* in its filamentous form inhibits infection of mice in a murine model (9–11). Histoplasmosis is usually self-limiting; however, immunocompromised individuals can develop a more severe form of disseminated histoplasmosis where the fungi infects other organs like the liver, kidneys, or spleen (12). A unique feature of *Hc* is its ability to become an intracellular pathogen of phagocytes, thus shielding it from the host immune system and providing an uninhibited vehicle for dissemination. *H. capsulatum* yeasts produce several virulence factors to evade killing and establish its niche within phagocytic cells (13–20). As such, understanding these virulence factors and their function is paramount to developing novel antifungal therapies to combat infection.

*DDR48* is a stress response protein shown to be important in combatting oxidative stress and antifungal drugs. No definitive function has been determined for *DDR48*; however, the protein contains multiple repeats of the peptide sequence Ser-Asn-Asn-X-Asp-Ser-Tyr-Gly, where X is either Asn or Asp that seem to be conserved between fungal species (21, 22). In *C. albicans*, *DDR48* is highly expressed during *in vivo* infections and is required for detoxification of the potent reactive oxygen species (ROS) hydrogen peroxide (21–24). A haploinsufficient *DDR48* mutant strain in *C. albicans* was also found to be more susceptible to killing by the common antifungals itraconazole, fluconazole, and ketoconazole when compared to a wild-type, *DDR48*-expressing strain (25–27). Hromatka et al. performed a genomic DNA microarray on *C. albicans* after exposure to nitric oxides for 10 minutes and found that *DDR48* was upregulated by 1.9-fold. These data demonstrate that *DDR48* is responsive to reactive nitrogen species (RNS) in *C. albicans*. They also found that *DDR48* was induced in a mutant strain devoid of nitric oxide dioxygenase (*YHB1*) activity (28). Another group found that expression of *DDR48* in response to amino acid starvation is dependent upon the amino acid biosynthesis transcriptional activator *GCN4*. They performed transcriptional profiling under amino acid starvation conditions in wildtype and a *gcn4Δ* mutant strain and found that *DDR48* expression was upregulated by 2.1-fold in the *gcn4Δ* mutant strain (29). The virulence transcription factor *CPH1*, which is involved in the formation of hyphae and psudohyphae has been shown to induce *DDR48* expression under hyphal inducing conditions (30). In contrast, in *C. albicans* strains devoid of the transcriptional repressors *TUP1* or *NRG1*, *DDR48* baseline transcriptional expression was overexpressed by roughly 15-fold (87). Nantel et al. performed microarray analyses of *C. albicans* cells in the presence of fetal bovine serum (FBS) at hyphal-inducing temperatures (37°C) and found that *DDR48* expression was upregulated by 2.9-fold (31). Another group repeated these experiments and also found that *DDR48* was upregulated in the presence of FBS as well as Spider’s medium, which promotes formation of hyphae (27). It was recently demonstrated that *DDR48* was highly induced in all phases of C. albicans infection in vivo and isolated *DDR48* protein in an extract of cell wall immunogenic proteins (32). This would suggest that *DDR48* is found in the cell wall in C. albicans, a characteristic unique to *C. albicans DDR48*. Kusch et al. performed a Coomassie stain on a 2D gel from a stationary growth phase culture of *C. albicans* and found *DDR48* to be one of the 50 most highly abundant proteins during stationary growth phase (33). Banerjee and associates performed a genome-wide steroid response study on *C. albicans* and found *DDR48* induced by 2-fold 30 minutes after the addition of 1mM progesterone to yeast cells. They also demonstrated that *DDR48* is among a unique subset of genes that are induced by the presence of ketoconazole, amphotericin b, and 5-fluorocytosine, as well as progesterone (34). Cleary et al. confirmed that *DDR48* is involved in the flocculation response as well as resistance to a variety of cellular stressors. The DDR48 mutant was more susceptible to 4NQO and amphotericin b treatment, demonstrating that *DDR48* is required for stress response and antifungal drug resistance in *C. albicans* (25). Another study performed in 2014 expounded on the implications of *DDR48’s* role in antifungal drug resistance. They found that clinical isolates of *C. albicans* that were resistant to fluconazole had significantly increased levels of *DDR48* mRNAs than those isolates that were fluconazole-sensitive (24). Based on these observations in *C. albicans*, we asked if *DDR48* is required for pathogenesis of *H. capsulatum*. In this study, we aimed to elucidate the role of *DDR48* in *H. capsulatum* by generating a *DDR48*-deficient strain and subjecting it to a battery of stressors and antifungal agents. We have demonstrated that a *DDR48*-deficient mutant is more susceptible to the ROS generators hydrogen peroxide and paraquat and more sensitive to the antifungal drugs ketoconazole and amphotericin B. We have also shown that the loss of *DDR48* results in a substantial decrease in *H. capsulatum* survival within macrophages. This study provides convincing results that *DDR48* is a suitable candidate to investigate as a potential antifungal therapy to make *H. capsulatum* more susceptible to killing by the host immune system.

## Results

### Identification of *DDR48* in *H. capsulatum*

*H. capsulatum DDR48* was originally isolated in our lab from a subtractive cDNA hybridization library enriched to identify transcripts whose expression was up-regulated in *Histoplasma* mycelia compared to *Histoplasma* yeasts. Transcripts identified in this library were named as “mold-specific” plus the number corresponding to when it was identified chronologically (e.g. *mold-specific 8; MS8*). Using this naming system, *DDR48* was originally referred to as *mold-specific 95* (*MS95*) (35). We performed a reciprocal protein BLAST of *MS95* using the *Saccharomyces* Genome Database (SGD) and found it to be an orthologue of the *S. cerevisiae DDR48* gene, sharing 48.6% identity and a corresponding E-value of 7.0 × 10^−22^ (**Table S2**). For continuity between fungal species in the literature, we updated the name from *MS95* to *DDR48*. *HcDDR48* encodes a 314 amino acid protein containing a total of 9 SNN(N/D)DSYG repeats, consistent with *DDR48* in other fungal species. Using the NCBI Conserved Protein Domain Family tool, we found *Histoplasma DDR48* to contain a conserved PTZ00110 (CDD) helicase domain belonging to the CL36512 domain superfamily. The PTZ00110 domain family consists of DEAD-box ATP-dependent RNA helicases involved in various aspects of RNA biosynthesis, maturation, and degradation (36, 37). To ascertain phylogeny of *Histoplasma DDR48*, we constructed a phylogenetic tree. We determined that *Histoplasma DDR48* is most closely related to *Paracoccidioides brasiliensis* and *Blastomyces dermatitidis* (**Figure 1**).

**Figure 1:**
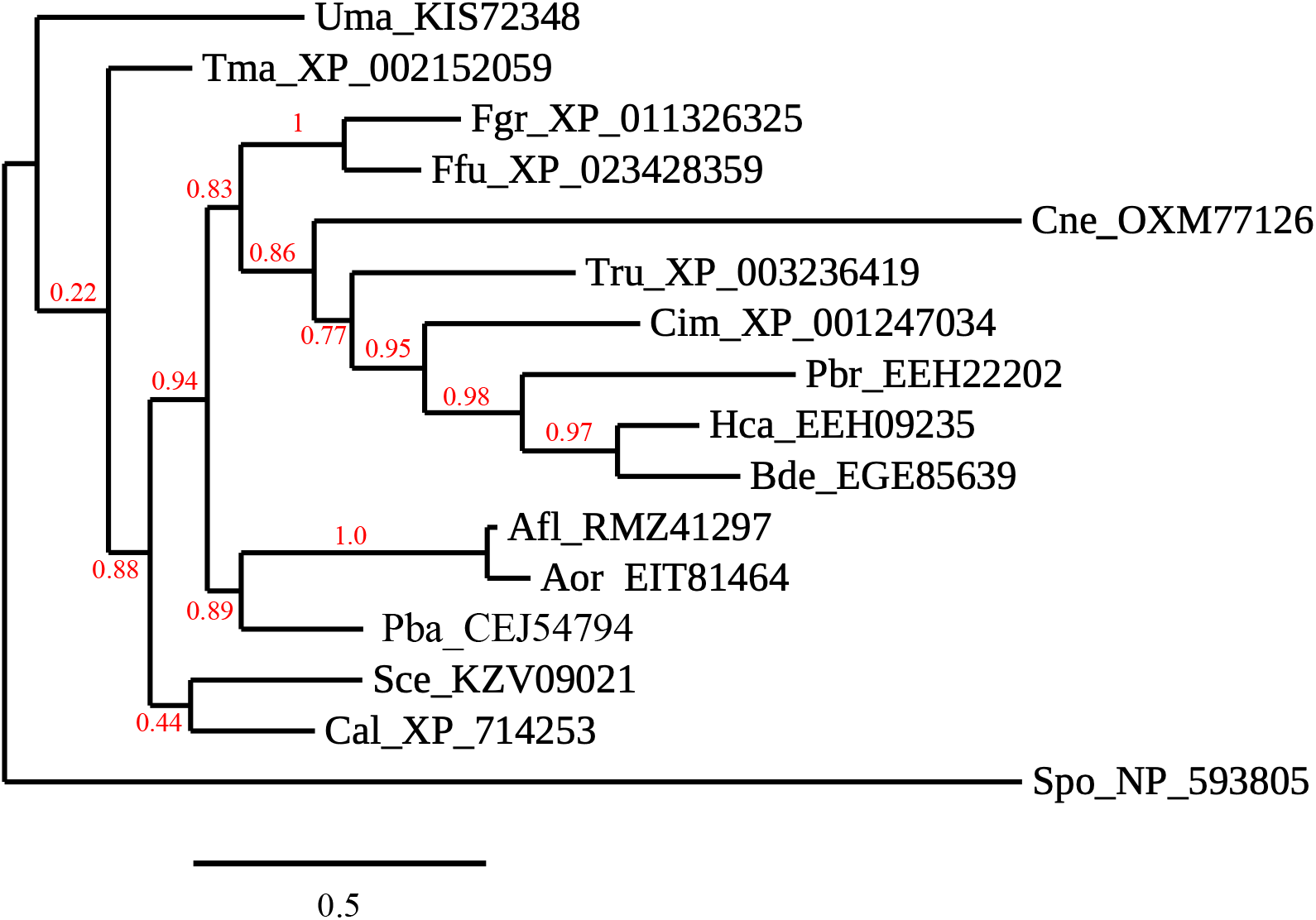
Phylogeny of *DDR48*. Phylogenetic analysis showing the relationship between *Histoplasma DDR48* and the gene in other fungi. Accession numbers are given for proteins from *Ustilago maydis (Uma)*, *Talaromyces marneffei (Tma)*, *Fusarium graminearum (Fgr)*, *Fusarium fujikuroi (Fju)*, *Cryptococcus neoformans var grubii (Cne)*, *Trichophyton rubrum (Tru)*, *Coccidioides immitis (Cim)*, *Paracoccidioides brasiliensis (Pbr)*, *Histoplasma capsulatum (Hca)*, *Blastomyces dermatitidis (Bde)*, *Aspergillus flavus (Afl)*, *Aspergillus oryzae (Aor)*, *Penicillium brasilianum (Pba)*, *Saccharomyces cerevisiae (Sce)*, *Candida albicans (Cal)*, and *Schizosaccharomyces pombe (Spo)*.

To confirm the results of the cDNA library that *DDR48* transcript levels are enriched in *Histoplasma* mold under optimal growth conditions, we performed quantitative, real-time PCR (qRT-PCR) and northern blot analysis on *Histoplasma* mold and *Histoplasma* yeasts from two laboratory strain, G186AR and G217B, to quantify *DDR48* expression. We determined that *DDR48* is expressed 6-fold higher in *Histoplasma* mold versus *Histoplasma* yeasts, compared to the constitutively expressed *Histone H3* gene (*HHT1*) in both strains of *H. capsulatum* when grown in the rich HMM broth (**Figure 2A**).

**Figure 2:**
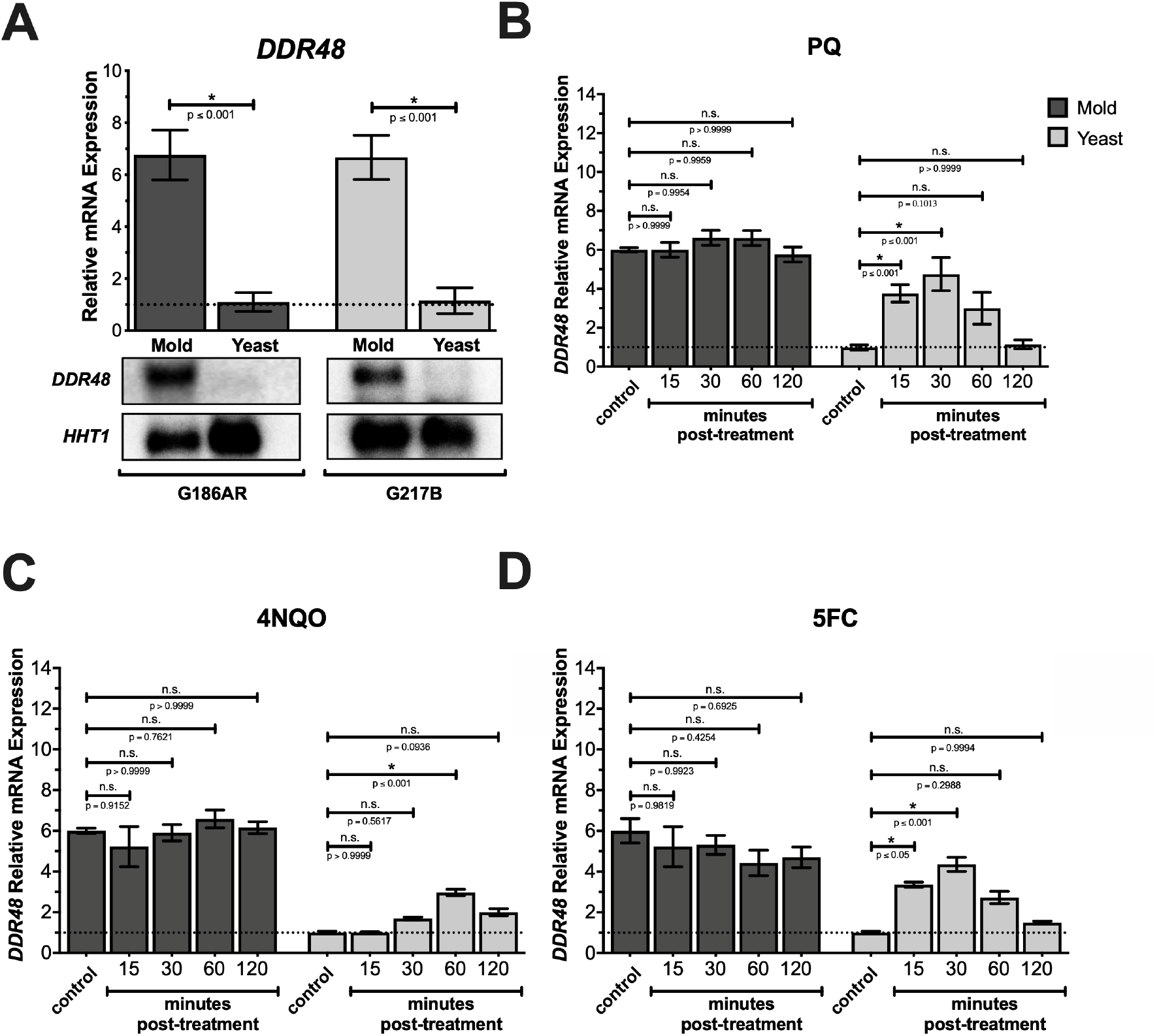
*DDR48* is constitutively expressed in mold-phase *Histoplasma* and inducible by stress in yeast-phase *Histoplasma*. **(A)** qRT-PCR and northern blot were performed on mold-phase and yeast-phase *Hc* samples from two clinically relevant strains, G186AR and G217B. qRT-PCRs were performed on mold-phase and yeast-phase wild-type (*DDR48 (+)*) cells under optimal conditions (control) and 15, 30, 60, and 120 minutes after addition of **(B)** paraquat (PQ), **(C)** 4-nitroquinolone-1-oxide (4NQO), and **(D)** 5-fluorocytosine (5FC). *HHT1* was used as a normalizer gene utilizing the *ΔΔ*_*ct*_ method to normalize mRNA levels to the no treatment control (dashed line).

### *Histoplasma DDR48* gene expression is responsive to Oxidative Stress Agents and DNA-Damaging Agents

Since *DDR48* has been characterized as a stress-response element in other fungal pathogens, we sought to determine if *DDR48* is involved in oxidative stress response in *H. capsulatum*. We first determined if the addition of oxidative stress led to any detectable changes in *DDR48* gene expression. This was accomplished by analyzing *DDR48* mRNA levels before and after the addition of the superoxide generator paraquat. We found that *DDR48* expression increased (4.5-fold) within 15 minutes post-paraquat addition with *DDR48* expression peaking at 30 minutes (**Figure 2B**). Within 120 minutes after exposure, *DDR48* mRNA levels returned to basal levels. Interestingly, upregulation of *DDR48* was only observed in *Histoplasma* yeasts, as there were no detectable changes in *DDR48* expression up to 120 minutes after the addition of paraquat (**Figure 2B**). These results were not surprising given our data above showed that *DDR48* expression in *Histoplasma* mold is 6-fold higher that *Histoplasma* yeasts to begin with (**Figure 2A**). These data propose that *DDR48* is constitutively expressed in mold-phase *H. capsulatum*; whereas, *DDR48* expression is inducible in *Histoplasma* yeasts when exposed to oxidative stress. We repeated these same experiments except we supplemented 4-nitroquinoline-1-oxide (4NQO) (**Figure 2C**) or 5-fluorocytosine (**Figure 2D**), a DNA damaging agent and DNA/RNA biosynthesis inhibitor, respectively. Results from these experiments indicate that *DDR48* expression increases in response to DNA damage as well as oxidative stress.

### Loss of *DDR48* Function Increases Sensitivity to Oxidative Stress

Since we have demonstrated that *DDR48* expression is responsive to oxidative stress additions, we determined if there are any differences in growth between a wild-type and *ddr48Δ* strain when grown in the presence of oxidative stress. We performed a series of growth curves where mid-log cells were cultured in rich HMM medium and supplemented with various concentrations of paraquat or hydrogen peroxide (H_2_O_2_). We included H_2_O_2_ as well since superoxides generated by paraquat are readily broken down to peroxides, thus *Histoplasma* cells will be challenged with both ROS *in vivo* (38). After 48 hours of incubation, turbidity values were recorded and normalized to no treatment controls. A significant decrease in growth of *ddr48Δ* yeasts was observed in those samples supplemented with paraquat and H_2_O_2_ as compared to wildtype controls (**Figure 3A**). Growth of the complemented *ddr48Δ + DDR48* strain was restored to levels observed in the wild-type samples (**Figure 3B**).

**Figure 3:**
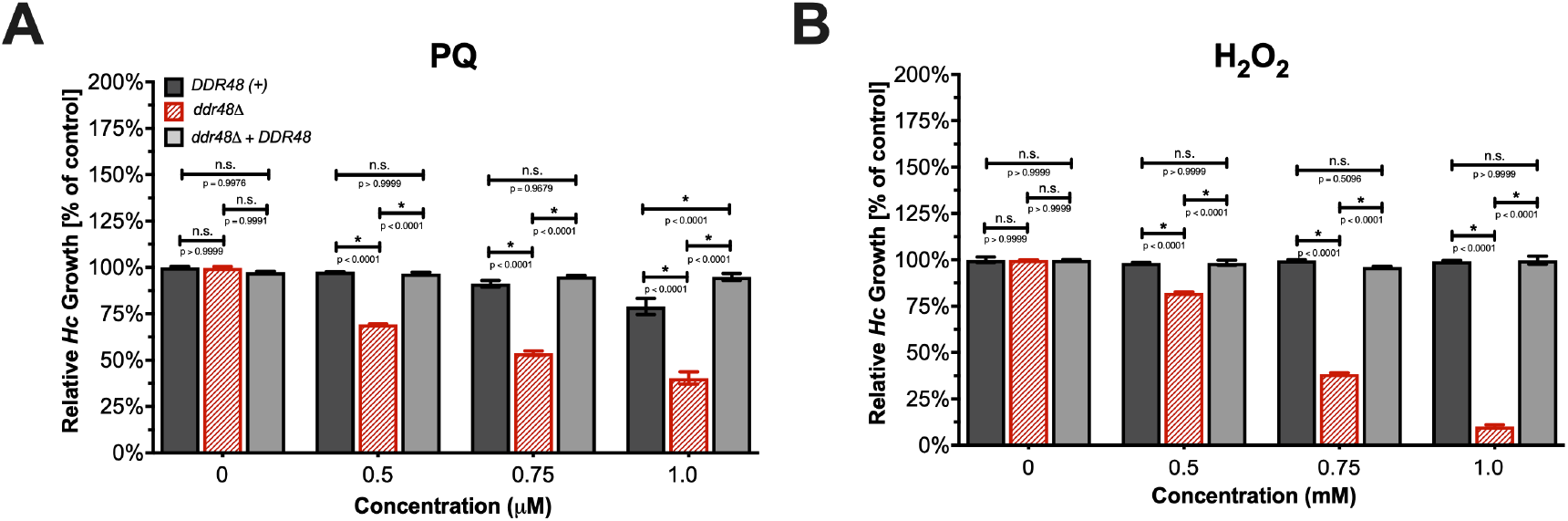
*DDR48* protects *Histoplasma* yeasts against reactive oxygen species. Survival of *DDR48 (+)* yeasts, *ddr48Δ* yeasts, and *ddr48Δ + DDR48* yeasts after 4-days of growth in liquid HMM broth supplemented with various concentrations of **(A)** paraquat (PQ) or **(B)** hydrogen peroxide (H_2_O_2_). Percent survival was normalized to *DDR48 (+)* no treatment control growth levels.

### Loss of *DDR48* Alters Cytosolic Catalases and Superoxide Dismutase Gene Expression

Since the loss of *DDR48* yielded a severe growth deficit when cells were under oxidative stress, we next performed gene expression analysis on *Histoplasma* catalases (*CatA, CatB,* and *CatP*) and superoxide dismutases (*SOD1* and *SOD3*) before and after exposure to paraquat. In wildtype *Histoplasma* yeasts, *CatA, CatB, CatP,* and *SOD1* gene expression increases after exposure to paraquat for 30 minutes. There were no detectable differences in *CatB* or *SOD3* (data not shown) between wildtype and *ddr48Δ* yeasts (**Figure 4C**). Interestingly, even before paraquat treatment *CatA, CatP,* and *SOD1* gene expression was decreased to almost undetectable levels in *ddr48Δ* yeasts (**Figure 4A,4B,4D**). Once treated with paraquat, gene expression levels of *CatA, CatP,* and *SOD1* increased, albeit by the same fold-change but not the same magnitude since they were already much lower than wildtype levels before treatment. It is worth noting that changes in *CatB* and *SOD3* gene expression was not surprising since *CatB* and *SOD3* are extracellular enzymes.

**Figure 4:**
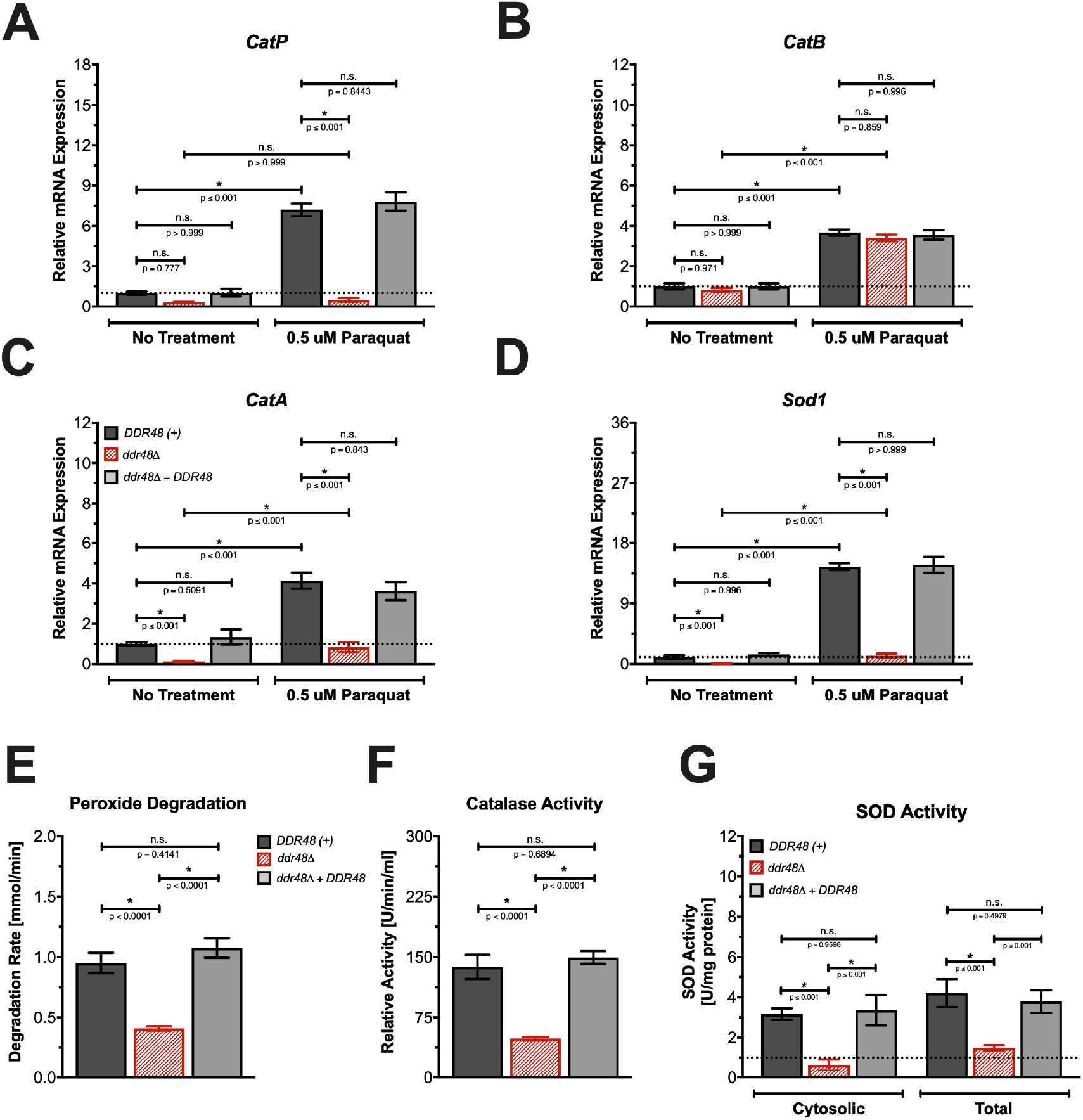
*DDR48* modulates *Histoplasma* catalases and superoxide dismutases in response to oxidative stress. qRT-PCRs were performed on *DDR48 (+),ddr48Δ,* and *ddr48Δ + DDR48* yeasts under optimal conditions (control) and 30 minutes after the addition of 0.5 μM Paraquat to determine gene expression levels of **(A)** *CatP*, **(B)** *CatB*, **(C)** *CatA*, and **(D)** *SOD1*. *HHT1* was used as a normalizer gene utilizing the ΔΔ_ct_ method to normalize mRNA levels to the no paraquat control (dashed line). **(E)** Hydrogen peroxide degradation rate, **(F)** relative catalase protein activity, and **(G)**(SOD) enzymatic activity were measured in *DDR48 (+),ddr48Δ,* and *ddr48Δ + DDR48 Hc* strains.

To confirm that the changes in catalase and superoxide dismutase gene expression seen in a *ddr48* mutant corresponds with actual protein levels, we measured catalase and superoxide dismutase (SOD) enzymatic activity in wildtype, *ddr48Δ*, and *ddr48Δ + DDR48* yeasts. We determined that the rate of hydrogen peroxide destruction was decreased by 57% in *ddr48Δ* yeasts (**Figure 4E**). H_2_O_2_ degradation rate was brought back to near wild-type levels in *ddr48Δ + DDR48* yeasts as expected. Furthermore, enzymatic activity of total catalases was determined for each strain to confirm that the decrease in degradation rate of H2O2 was due to a decrease in catalase activity. Catalase activity of *ddr48Δ* yeasts was 64% less than activity of the wildtype controls (**Figure 4F**). Complementation of *DDR48* restored catalase activity to wildtype levels (FIGURE). Next, we calculated cytosolic SOD activity and total SOD activity for wildtype yeasts, *ddr48Δ* yeasts, and *ddr48Δ + DDR48* yeasts. Cytosolic SOD activity of *ddr48Δ* yeasts was determined to be 0.64 U/mg protein, which is an 80% decrease compared to wildtype levels (**Figure 4G**). Total SOD activity of *ddr48Δ* yeasts was also decreased by 64%; whereas, cytosolic and total SOD activity was restored to near wildtype levels in *ddr48Δ + DDR48* yeasts (**Figure 4G**). These results confirm that the loss of *DDR48* decreases the amount of catalase and superoxide dismutase enzymatic activity in optimal conditions.

### *DDR48* depleted *Histoplasma* have increased levels of glutathione-dependent redox transcripts

Since *ddr48Δ* yeasts are able to proliferate, albeit, at much lower rates, under oxidative stress, we examined other components of the oxidative stress response, specifically the glutathione-dependent redox system. We performed qRT-PCRs on the cytosolic thioredoxin reductase *TRR1* (**Figure 5C**), the cytosolic thioredoxin *TRX1* (**Figure 5D**), the glutamate-cysteine ligase *GSH1* (**Figure 5A**), and the glutathione synthetase *GSH2* (**Figure 5B**) in optimal growth conditions (HMM) and oxidative stress (HMM + paraquat) on all three strains. In wildtype yeasts there were no significant changes in mRNA levels of *HcTRR1, HcTRX1, HcGSH1,* and *HcGSH2* upon challenge with oxidative stress. Surprisingly, *HcTRR1, HcTRX1, HcGSH1,* and *HcGSH2* were all significantly upregulated in *ddr48Δ* yeasts challenged with oxidative stress. Transcript levels in *ddr48Δ + DDR48* complemented strain matched those seen in wildtype yeasts, as expected. These results suggest that the glutathione-dependent oxidative stress system is compensating for the loss of catalase and superoxide dismutase activity see in *ddr48Δ* yeasts.

**Figure 5:**
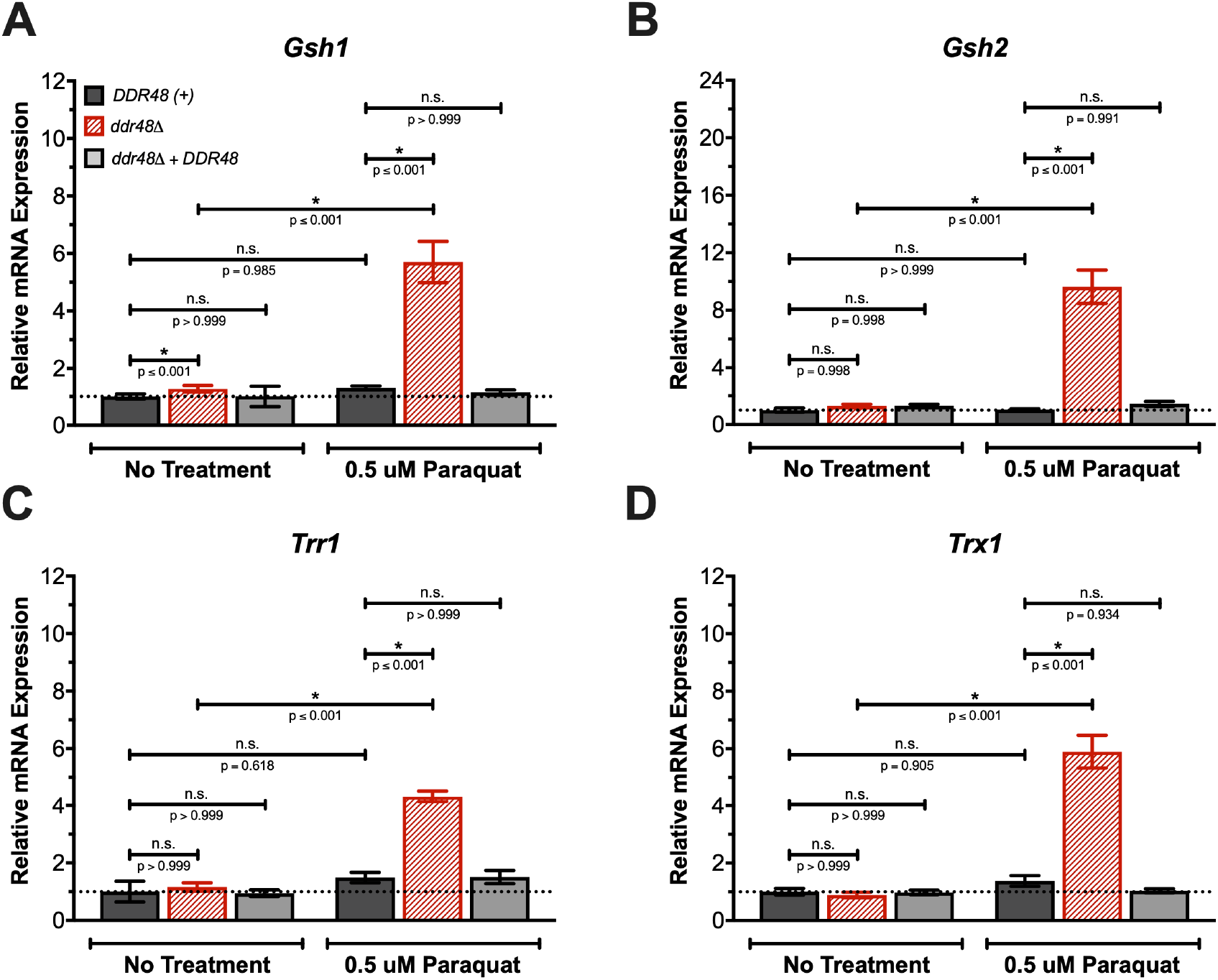
Endogenous glutathione machinery compensates for the loss of *DDR48* when subjected to ROS. qRT-PCRs were performed on *DDR48 (+),ddr48Δ,* and *ddr48Δ + DDR48* yeasts under optimal conditions (control) and 30 minutes after the addition of 0.5 μM paraquat to determine gene expression levels of **(A)** *GSH1*, **(B)** *GSH2*, **(C)** *TRR1*, and **(D)** *TRX1*. *HHT1* was used as a normalizer gene utilizing the ΔΔ_ct_ method to normalize mRNA levels to the no paraquat control (dashed line).

### *DDR48* depleted *Histoplasma* yeasts possess increased sensitivity to ketoconazole and amphotericin b

Increased *DDR48* expression has been linked to antifungal drug resistance in clinical isolates of the pathogenic fungi *C. alicans* (24). To determine if *HcDDR48* transcription levels are responsive to antifungal drug exposure, we first performed gene expression analysis on *DDR48 (+)* yeasts before and after exposure to amphotericin b and ketoconazole, two representatives of the common antifungal groups used in the treatment of invasive mycotic infections (39). We determined that *HcDDR48* mRNA accumulates after exposure to ketoconazole (**Figure 6A**) and amphotericin b (**Figure 6B**). To determine the sensitivity of *DDR48-*expressing and *DDR48*-depleted yeasts to these antifungals, we performed a microtiter plate assay to determine the minimum inhibitory concentration (MIC) and 50% inhibition (IC_50_) values. We determined that MIC values for ketoconazole (**Figure 6C**) and amphotericin b (**Figure 6D**) were significantly decreased by roughly 50% in *ddr48Δ* yeasts. The decrease in MIC values were accompanied by a substantial decrease in IC_50_ concentrations for both drugs in *ddr48Δ* yeasts, as expected. These data indicate that the loss of *DDR48* considerably decreases tolerance of *Histoplasma* yeasts to ketoconazole and amphotericin b.

**Figure 6:**
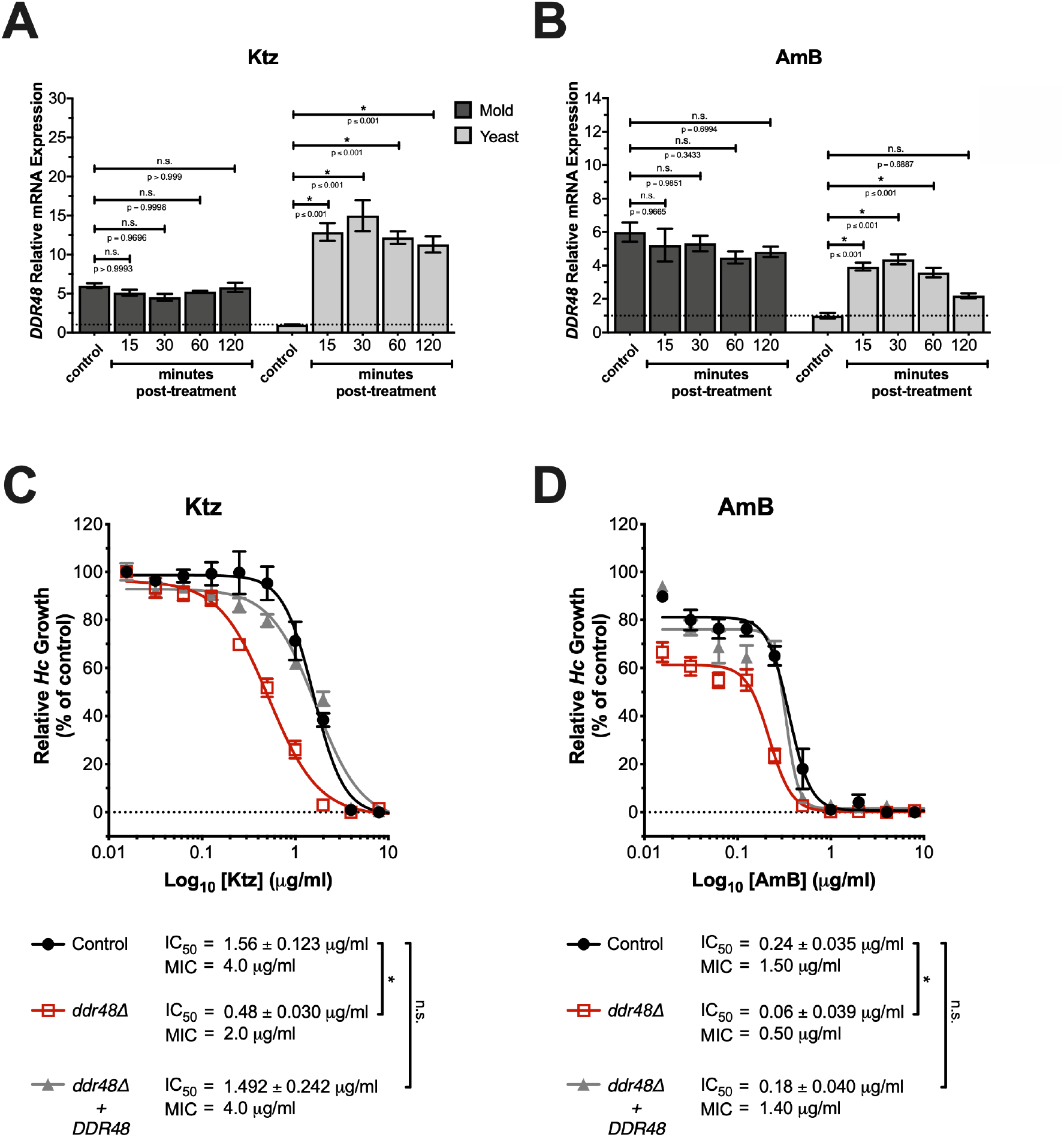
*DDR48* contributes to *Histoplasma* yeast’s resistance to ketoconazole and amphotericin-b. qRT-PCRs were performed on mold and yeast-phase wild-type (*DDR48 (+)*) cells under optimal conditions (control) and 15, 30, 60, and 120 minutes after addition of ketoconazole (Ktz) **(A)** or amphotericin b (AmB) **(B)**. *HHT1* was used as a normalizer gene utilizing the ΔΔ_ct_ method to normalize mRNA levels to the no treatment control (dashed line). Dose response curves were generated using ketoconazole (Ktz) **(C)** or amphotericin b (AmB) **(D)** using *DDR48 (+),ddr48Δ*, and *ddr48Δ + DDR48* yeasts. IC_50_ values were determined by non-linear regression and MIC values were determined by minimum drug concentration that yielded no *Hc* growth. IC_50_ and MIC values are represented as mean ± standard error of the mean (SEM).

### *DDR48* Depleted *Histoplasma* Contain An Aberrant Ergosterol Biosynthesis Pathway

Since amphotericin b and ketoconazole affect the ergosterol biosynthesis pathway, we performed gene expression analysis on the various components of the ergosterol biosynthesis pathway in before and after amphotericin b treatment. Interestingly, even before addition of amphotericin b *HcERG6* (**Figure 7A**) and *HcERG24* (**Figure 7B**) were significantly decreased in *ddr48Δ* yeasts. In contrast, *HcERG11a* (**Figure 7C**) and *HcERG11b* (**Figure 7D**) were significantly increased in *ddr48Δ* yeasts before amphotericin b challenge. Not surprisingly, transcript levels of all ergosterol biosynthesis gene tested increased upon exposure to amphotericin b in wildtype yeasts (**Figure 7E**). However, in *ddr48Δ* yeasts, there were no detectable changes in transcript levels of *HcERG3, HcERG4, HcERG5, HcERG6, HcERG7,* and *HcERG26* after amphotericin b exposure. Interestingly, levels of *HcERG24, HcERG25,* and *HcSRB1* increased in *ddr48Δ* yeasts, just not to the levels measured in wildtype data. *HcERG11a* levels increased beyond wildtype levels while *HcERG11b* levels matched those of wildtype yeasts after drug challenge (**Figure 7E**). Complementation of *HcDDR48* restored levels of each gene to those observed in wildtype yeasts. The data above demonstrate that the loss of *DDR48* disrupts the ergosterol biosynthesis pathway.

**Figure 7:**
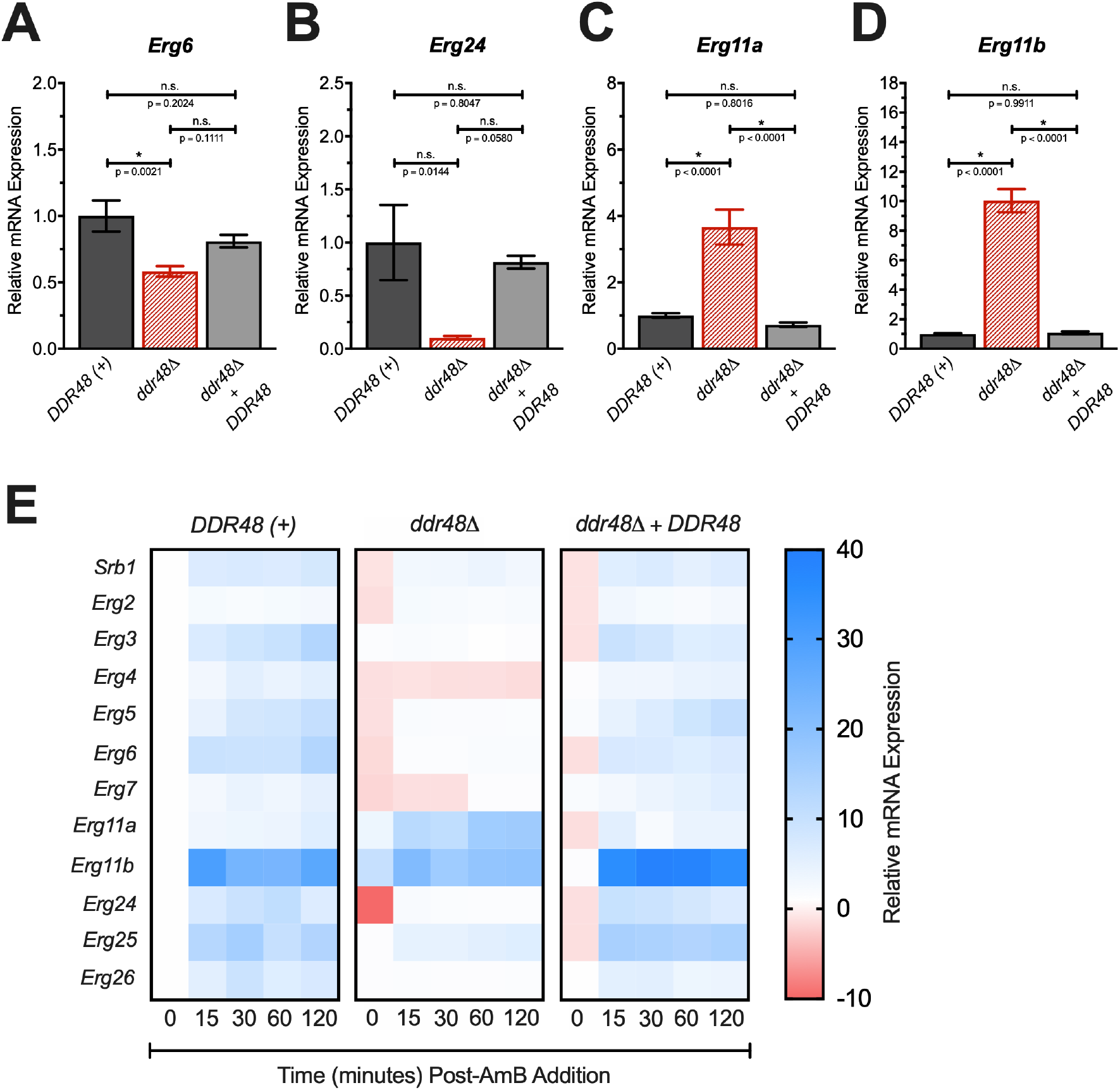
The loss of *DDR48* results in an aberrant ergosterol biosynthesis pathway. qRT-PCRs were performed on *DDR48 (+),ddr48Δ,* and *ddr48Δ + DDR48* yeasts under optimal conditions (HMM) to measure gene expression of **(A)** *Erg6,* **(B)** *Erg24,* **(C)** *Erg11a*, and **(D)** *Erg11b*. **(E)** Heat map depicting qRT-PCRs performed on *DDR48 (+),ddr48Δ,* and *ddr48Δ + DDR48* yeasts under optimal conditions (control) and 15, 30, 60, and 120 minutes after addition of amphotericin b. *Histone H3* was used as a normalizer gene utilizing the ΔΔ_ct_ method to normalize mRNA levels to the no treatment control.

### Optimal Survival Of *Histoplasma* Yeasts In Macrophages Is Dependent Upon A Functional *DDR48* Gene

Macrophages respond to *H. capsulatum* infection by generating reactive oxygen species (ROS) and the loss of *DDR48* results in decreased oxidative stress response by *Histoplasma* yeasts, we asked if *DDR48* depleted cells demonstrate altered intracellular survival within resting and IFNγ-activated murine macrophages. We first determined *DDR48* gene expression within macrophages. We found that within 4 hours post-infection *HcDDR48* transcript levels dramatically increased and remained elevated up to 24 hours post infection (**Figure 8A**). Next, we infected resting and IFNγ-activated murine macrophages with either *DDR48*-expressing or *DDR48*-depleted yeasts at various concentrations. Survival of *ddr48Δ* yeasts was decreased by roughly 50% in resting and IFNγ-activated macrophages in all assays tested (**Figure 8B,8C,8D**). Survival of the complemented *DDR48* strain was nearly identical to wildtype levels in each scenario.

**Figure 8:**
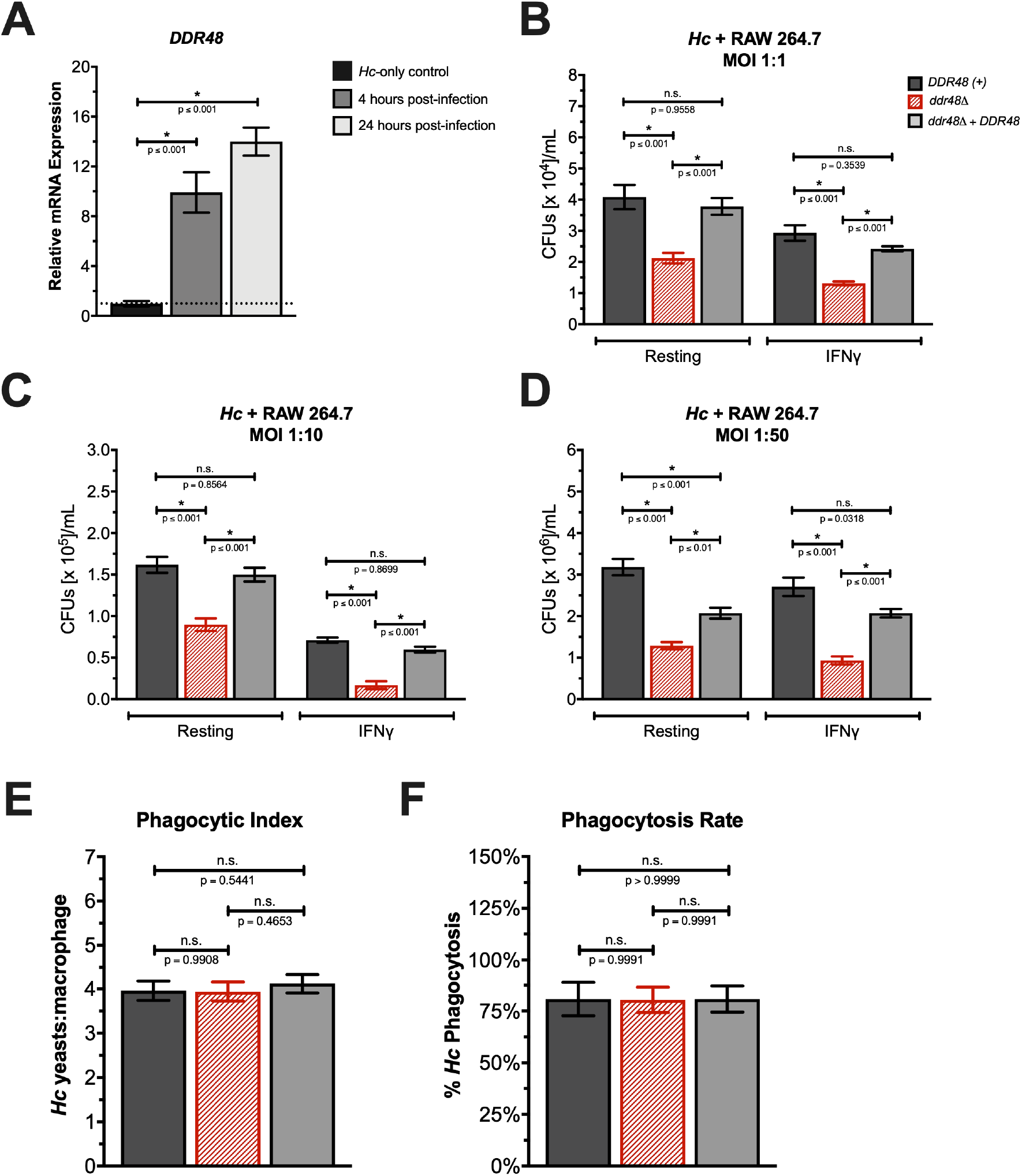
*DDR48* is upregulated upon uptake by macrophages and the loss of *DDR48* results in decreased *Histoplasma* survival in resting and IFNγ activated macrophages. **(A)** qRT-PCRs were performed on *DDR48 (+)* yeasts under optimal growth conditions (control), 4 hours post-infection, and 24 hours post-infection. *Histone H3* was used as a normalizer gene utilizing the ΔΔ_ct_ method to normalize mRNA levels to uninfected control. Infection of resting and activated murine macrophages with *DDR48 (+),ddr48Δ, and ddr48Δ + DDR48* were performed at a multiplicity of infection (MOI) or **(B)** 1:1**, (C)** 1:10, and **(D)** 1:50. We also determined the **(E)** phagocytic index and **(F)** phagocytosis rate for *DDR48 (+), ddr48Δ,* and *ddr48Δ + DDR48* yeasts using resting murine macrophages at an MOI of 1:5.

To ensure that the decrease in recovery of *ddr48Δ* yeasts is due to decreased survival within macrophages and not simply the result of different rates of phagocytosis, we determined if the same amount of *Hc* yeasts were being phagocytized between each strain as well as if the average number of macrophages taking up *Hc* yeasts was equal between the *Hc* strains being tested. To determine the average number of *Hc* yeasts being phagocytosed per macrophage, we performed a phagocytic index assay. We found no significant differences in the phagocytic index of wildtype yeasts, *ddr48Δ* yeasts, or *ddr48Δ + DDR48* yeasts (**Figure 8E**). There were no significant differences in the number of macrophages actively phagocytizing *Hc* between wildtype yeasts, *ddr48Δ* yeasts, or *ddr48Δ + DDR48* yeasts with roughly 80% of macrophages containing *Hc* yeasts in each trial (**Figure 8F**). These results show that the decrease in recovery of *ddr48Δ* yeasts was due to a decreased fitness of *ddr48Δ* yeasts within macrophages and not due to an entry defect. These data suggest that *DDR48* is potentially involved in pathogenesis of *Histoplasma* yeasts since a loss of *DDR48* function results in decreased *Histoplasma* survival within macrophages.

## Discussion

In this study, we aimed to increase our understanding of the *DDR48* gene in *Histoplasma capsulatum* and determine if *DDR48* is essential for response to oxidative stress, response to antifungal drugs, and survival within phagocytes. We employed a bioinformatics analysis and identified nine repeats of the amino acid sequence SNN(N/D)DSYG within the *Histoplasma DDR48* amino acid sequence, which is consistent with sequences of *DDR48* from the well characterized fungi *S. cerevisiae* and *C. albicans*. In an attempt to determine the function of *DDR48* in *Histoplasma*, we analyzed the amino acid sequence for conserved domains using NCBI’s Domain Architecture Retrieval Tool (DART) and found that the *Histoplasma DDR48* amino acid sequence contains conserved domains most closely related to a group of ATP-dependent RNA helicases (PTZ00110). This group of RNA helicases use ATP molecules to achieve their RNA un-winding activities hence they also exhibit ATP hydrolysis (40). A study on *DDR48* in *S. cerevisiae* concluded that *DDR48* possesses ATP and GTP hydrolysis abilities (21). This part of the study is purely theoretical modeling and algorithms; therefore, more direct analysis of *DDR48*’s function needs to be completed before we can determine *HcDDR48* function.

Most genes that are enriched in one growth phase of *Histoplasma* are essential for survival or pathogenesis. For example, the extracellular superoxide dismutase *SOD3* is only expressed in yeast-phase *Histoplasma* as extracellular oxidative stress defense is most likely to be needed when phagocytes undergo oxidative bursts as an effort to eliminate *Histoplasma* yeasts *in vivo* (20). If the gene in question is essential for maintaining a specific morphotype (mold or yeast), then the absence of the gene results in *Histoplasma* cells that are “locked” in one phase of growth, independent of temperature or transcriptional growth programs. An example would be the *Histoplasma MSB2* gene, which was found to be essential for formation of hyphae, as deletion of *MSB2* resulted in the cells being yeast-locked, where *Histoplasma* grew as yeasts at room temperature as well as 37°C (41).We have shown that *Histoplasma DDR48* is enriched in the mold-phase of growth in optimal conditions, which suggests that *DDR48* has an essential role that is specific to the mold-phase of growth. We showed this was not the case with *DDR48*, as mold-phase growth of the *ddr48Δ* mutant was phenotypically no different than *DDR48*-expressing strains, where both grew as hyphae at room temperature and yeasts at 37°C. Even though *DDR48* expression is enriched in mold-phase in optimal conditions, our gene expression studies of *DDR48* showed that it’s expression increased in response to oxidative stress in *Histoplasma* yeasts. We have shown that *DDR48* is constitutively expressed in mold-phase *Histoplasma* and inducible in *Histoplasma* yeasts. This brings up many questions regarding the study of phase-specific genes in dimorphic fungi. In past studies, if a gene was found to only be transcribed in one growth phase it was labelled as a phase-specific gene. Here we show that much more care should be taken when characterizing potential phase-specific genes. The higher basal transcription rate of *DDR48* in *Histoplasma* mold could be attributed to an increased exposure to oxidative stress as compared to *Histoplasma* yeasts, although this is just speculation. Pathogenic *Histoplasma* yeasts are in a controlled environment within the host, where exogenous oxidative stress is expected, thus the production of the secreted *SOD3* superoxide dismutase. Environmental conditions encountered by *Histoplasma* mold are variable and dependent upon geographic location, nutrient availability, and weather conditions (42–45). By having *DDR48* constitutively expressed in mold-phase *Histoplasma*, the fungi may be better poised to deal with the unpredictable environments it will encounter, though more studies would be needed in order to support this concept.

Growth of the *ddr48Δ* strain was severely inhibited by the presence of oxidative stress generators, demonstrating that *DDR48* is required for response to oxidative stress in *Histoplasma* yeasts. Complementation of the *ddr48Δ* mutant with an episomal copy of *DDR48* fully rescued resistance to oxidative stress to levels seen in the wildtype strain. We have also shown through our gene expression data that transcription of *DDR48* mRNAs is responsive to the presence of oxidative stress. An up-regulation of *DDR48* mRNAs was seen within 15 minutes after exposure of *Histoplasma* yeasts to oxidative stress. Maximum *DDR48* expression is achieved 30 minutes after oxidative stress exposure, thereafter mRNA levels returned to baseline. These results are consistent with experiments performed in *S. cerevisiae* and *C. albicans* that determined *DDR48* is involved in the oxidative stress response (21, 26, 46–49). The cytosolic superoxide dismutase *SOD1* and the cytosolic catalase *CatP* are responsible for detoxification of intracellular ROS in *Histoplasma* yeasts (20, 50). Gene expression of *SOD1* and *CatP* increases when there is an increase in intracellular ROS to rapidly eliminate them. Interestingly, gene expression of both *SOD1* and *CatP* was significantly decreased in the *ddr48Δ* strain in liquid HMM cultures, even without oxidative stress. While *SOD1* gene expression was decreased in *ddr48Δ* yeasts, when challenged with the ROS generator paraquat, *SOD1* mRNA levels still increased but not to the same magnitude of wildtype *Histoplasma* yeasts. This suggests that *DDR48* is involved in basal regulation of *SOD1* transcription and not necessarily in its regulation in response to oxidative stress. On the other hand, *CatP* gene expression in *ddr48Δ* yeasts was not responsive to an increase in ROS, suggesting that *DDR48* is involved in modulating *CatP* in response to oxidative stress in addition to basal regulation of transcription. These findings suggest that *DDR48*-dependent modulation is different for each gene. We confirmed these results on the enzymatic level and determined that catalase and superoxide dismutase enzymatic activities were decreased in *ddr48Δ* cells and activity was rescued in the *DDR48* complementation strain, consistent with our transcription data. We conclude that *DDR48* directly or indirectly modulates intracellular catalase and superoxide dismutase activity since the lack of *DDR48* leads to a decrease in growth when exposed to oxidative stress. It is important to mention that *DDR48* is not required for detoxification of ROS generated by basal cellular metabolism, as deficits in growth are only seen when subjected to stress; although, this lack of phenotype could be as a result of other compensatory pathways that is sufficient enough to rescue growth under optimal conditions.

Though cytosolic ROS detoxification machinery is dysregulated in *ddr48* yeasts, the glutathione-dependent ROS detoxification machinery compensates. When *ddr48Δ* yeasts were challenged with ROS, the cytosolic thioredoxin reductase *TRR1*, the cytosolic thioredoxin *TRX1*, the glutamate-cysteine ligase *GSH1*, and the glutathione synthetase *GSH2,* were all transcriptionally up-regulated; whereas, when *Histoplasma* yeasts containing a functional copy of *DDR48* are challenged with ROS, there are no observable changes in gene expression of the glutathione-dependent transcripts *GSH1, GSH2, TRR1,* and *TRX1*. These results demonstrate that one way *Histoplasma* yeasts compensate for the loss of *DDR48* is by activating alternative ROS detoxification pathways to aid in survival. These results are consistent with studies in *S. cerevisiae* where yeasts lacking the cytosolic thioredoxin system were more susceptible to killing by oxidative stress (51).

The antifungals fluconazole and itraconazole, in combination with amphotericin B, are mainline treatments for infection with *Histoplasma capsulatum* (52). One of the leading causes of treatment failure in HIV-positive patients with *H. capsulatum* infections is resistance to azole antifungals, thus new treatment options are needed to decrease relapse prevalence (53). Here we have shown that *ddr48Δ* yeasts are more susceptible to killing by ketoconazole and amphotericin b. These results are consistent with studies in *C. albicans* where researchers found that fluconazole-resistant isolates of *C. albicans* from patients had higher expression levels of *DDR48*. They also found a high correlation of *DDR48* and azole resistant genes, suggesting *DDR48* is a candidate for potentially novel antifungal therapies (22). One way *DDDR48* seems to exert its protective effects is by modulating the ergosterol biosynthesis pathway, which is consistent with studies in *C. albicans* where a gain of function mutation leading to increased expression of ergosterol biosynthesis genes led to an increase in antifungal drug resistance (54).

We also demonstrated that *Histoplasma* survival within macrophages is dependent on a functional *DDR48* gene. We have shown that *Histoplasma* survival is consistently decreased by half in *ddr48Δ* yeasts at multiple *Histoplasma* doses. This could be due to the increased oxidative stress that *Histoplasma* encounters within phagocytes or that there is less entry of *ddr48Δ* yeasts into phagocytic cells. We demonstrated that the decrease in survival is not due to an entry defect as the same phagocytosis rates were observed in *DDR48*-expressing strains and mutant strains. When infected with fungal pathogens, phagocytes initiate an oxidative burst via the NADPH oxidase system that serves to control fungal growth (55). *Histoplasma* yeasts do contain an extracellular catalase, *CatB*, and an extracellular superoxide dismutase, *SOD3*, to counter the ROS derived from phagocytes. *CatB* gene expression was not altered by the loss of *DDR48*, suggesting that the decreased survival is not directly due to less extracellular enzyme activity. Since endogenous ROS is increased by the decreased cytosolic catalase and superoxide dismutase activity in *ddr48Δ* yeasts, the extracellular ROS detoxification machinery could be compensating for this loss and indirectly leading to more effective killing of *Histoplasma* within phagocytes. In these experiments, we used RAW 264.7 macrophage-like cells, a common macrophage cell line used in phagocytosis studies (56). Data from human macrophages might provide data more suited towards therapeutic value.

We have demonstrated that *DDR48* functions in various pathways in *H. capsulatum* to aid in fungal stress response and proliferation. We have shown that *DDR48* is involved in the oxidative stress response and that the loss of *DDR48* leads to increased sensitivity of *Histoplasma* yeasts to ROS. *DDR48* appears to modulate the transcription of *SOD1* and *CatP*, a cytosolic superoxide dismutase and catalase, respectively. We also demonstrated that *DDR48* is essential for survival of *Histoplasma* yeasts within resting and activating murine macrophages, suggesting it is required for pathogenesis. *In vivo* research will need to be conducted before we can conclude that *DDR48* is needed pathogenesis as *in vitro* data does not always correlate to an *in vivo* system, where the host immune system is involved (57, 58). We also concluded that the loss of *DDR48* led to increased sensitivity of *Histoplasma* yeasts to amphotericin B and ketoconazole, suggesting that *DDR48* could be exploited as a potential therapeutic target. All of these results taken together point to *DDR48* possessing a central role in modulating the fungal cellular stress response. We cannot exclude the possibility that *DDR48* is a global modulator of cellular stress response and therefore any stress encountered by the *ddr48Δ* strain will result in a difference in growth compared to wildtype *Histoplasma* yeasts. To fully understand which genes are *DDR48*-dependent a more global approach will need to be employed, such as RNA sequencing.

Interestingly, Jin and associates identified *DDR48* as a component of processing bodies (P-bodies) and glycolytic bodies (G-bodies) in *C. albicans* (59). P-bodies are mRNA granules whose function is in maintaining the balance between translating mRNAs and mRNA degradation. P-bodies contain aggregates of mRNA decay machinery and mRNAs destined for repression/degradation or balanced reentry into translation. In response to cellular stress, such as glucose starvation, stress granules are formed from P-bodies to aid in response and recovery from the stress (60–62). Each of these mRNA granules relies upon RNA binding proteins for proper structure and function (63, 64). Mechanisms by which *DDR48* could potentially be participating in formation and processing within these specialized non-membrane bound organelles could be by 1) chaperoning mRNAs into the bodies or 2) serving as a linker between RNAs within the P-body. We have constructed a hypothetical model to illustrate these possible functions of *DDR48*, which can be seen in **Figure 9**. It should be noted that this is a purely theoretical model constructed in order to direct further research into the function of *DDR48* in pathogenic fungi. We have demonstrated that *H. capsulatum* relies on *DDR48* to adapt and recover in response to cellular stress and confirmed that it is needed for optimal survival in macrophages, response to oxidative stress, and response to antifungal drugs. More research into the function of *DDR48* will no doubt uncover more ways in which it is responsible for survival against cellular stressors in fungal pathogens. Given the broad range of processes that are dependent upon *DDR48*, thought should be given to determining if it could be a successful therapeutic target for those predisposed or suffering from endemic fungal pathogens, as a pathogen that is not poised to adapt when exposed to various cellular stressors could tip the balance in favor of the host’s immune system and eventual elimination of the fungal pathogen.

**Figure 9:**
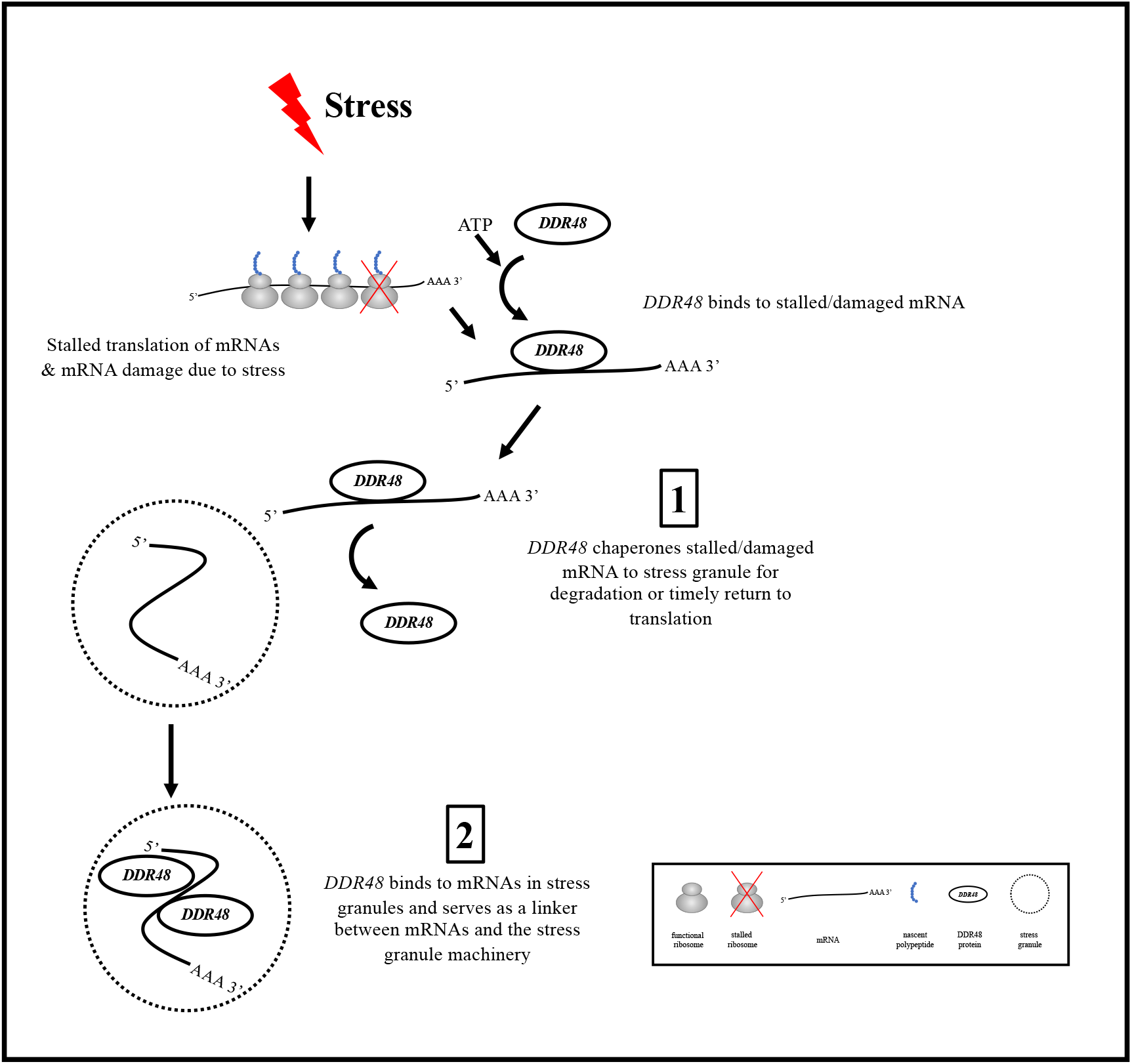
Hypothetical pathway depicting possible functions of *DDR48* in RNA binding in response to cellular stress. *DDR48* binds to stalled/damaged mRNAs and either **(1)** chaperones stalled/damaged mRNAs to stress granules for degradation/timely return to translation or **(2)** *DDR48* binds to mRNAs in stress granules and serves as a linker between mRNAs and the stress granule machinery; both scenarios result in resolution of endogenous stress in a *DDR48*-dependent manner.

## Supplemental Figure & Table Legends

**Figure S1:**
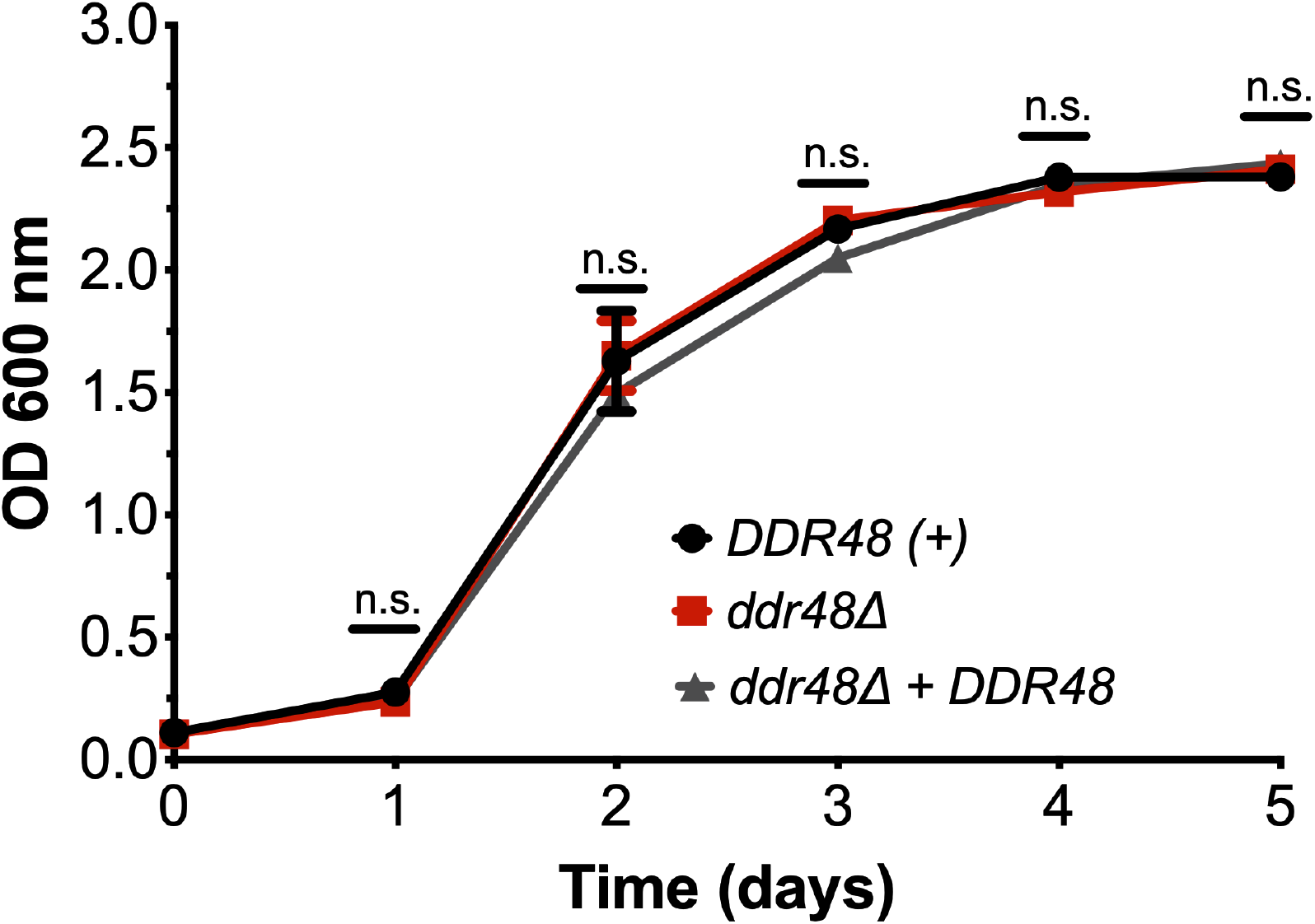
Growth curve of *Histoplasma* strains. Growth curve demonstrating no detectible differences in growth of wildtype, *ddr48Δ,* and *ddr48* + *DDR48* yeasts in liquid HMM growth media.

**Figure S2:**
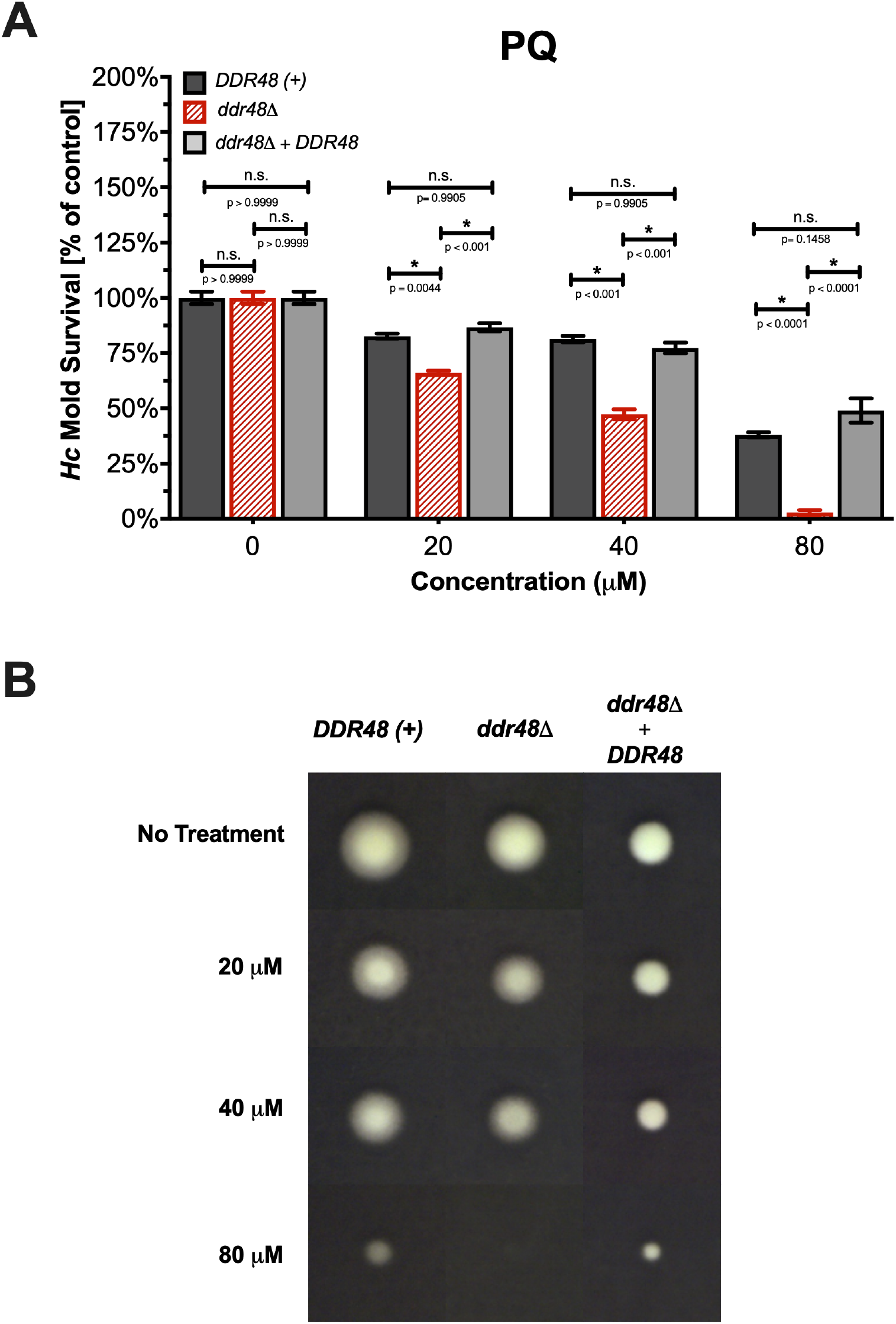
Loss of *DDR48* leads to increased susceptibility of mold-phase *Histoplasma*. Quantitative (A) and qualitative (B) growth of wildtype, *ddr48Δ,* and *ddr48 + DDR48 Hc* mold strains on various concentrations of paraquat

**Figure S3:**
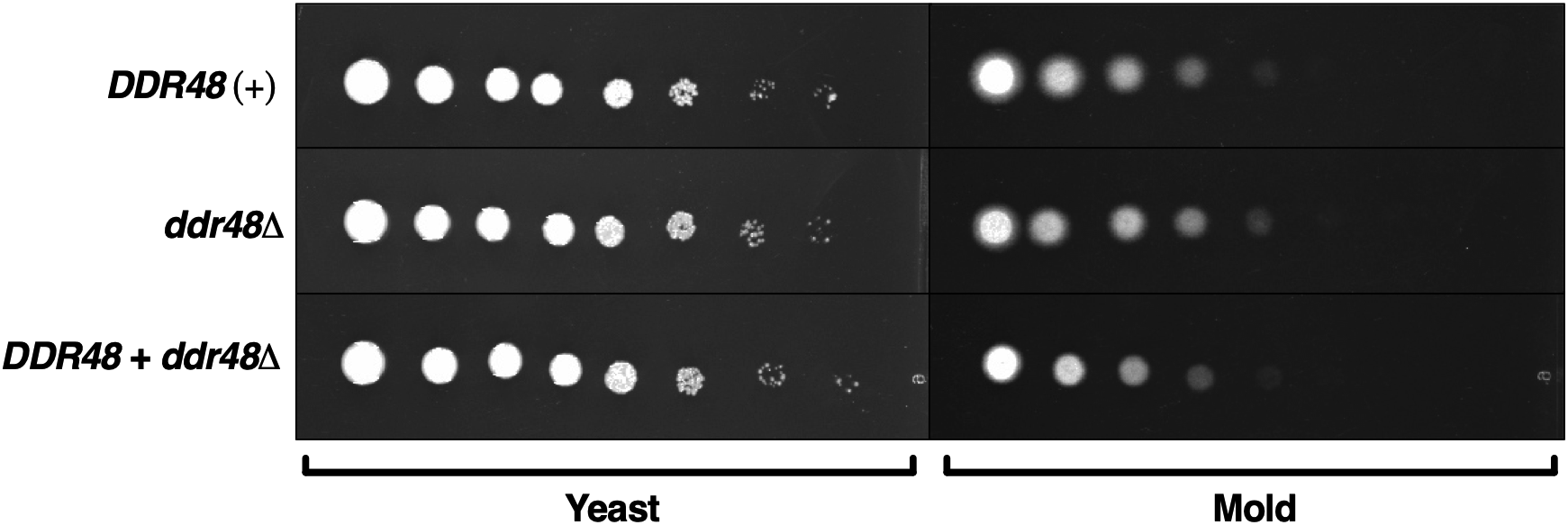
There are no detectable differences in growth of *DDR48(*+) and *ddr48Δ* yeasts on solid growth medium. Growth of wildtype yeasts and mold, *ddr48Δ* yeasts and mold, *and ddr48* + *DDR48* yeasts and mold on rich HMM medium depicting no qualitative changes in growth rate under optimal growth conditions.

**Figure S4:**
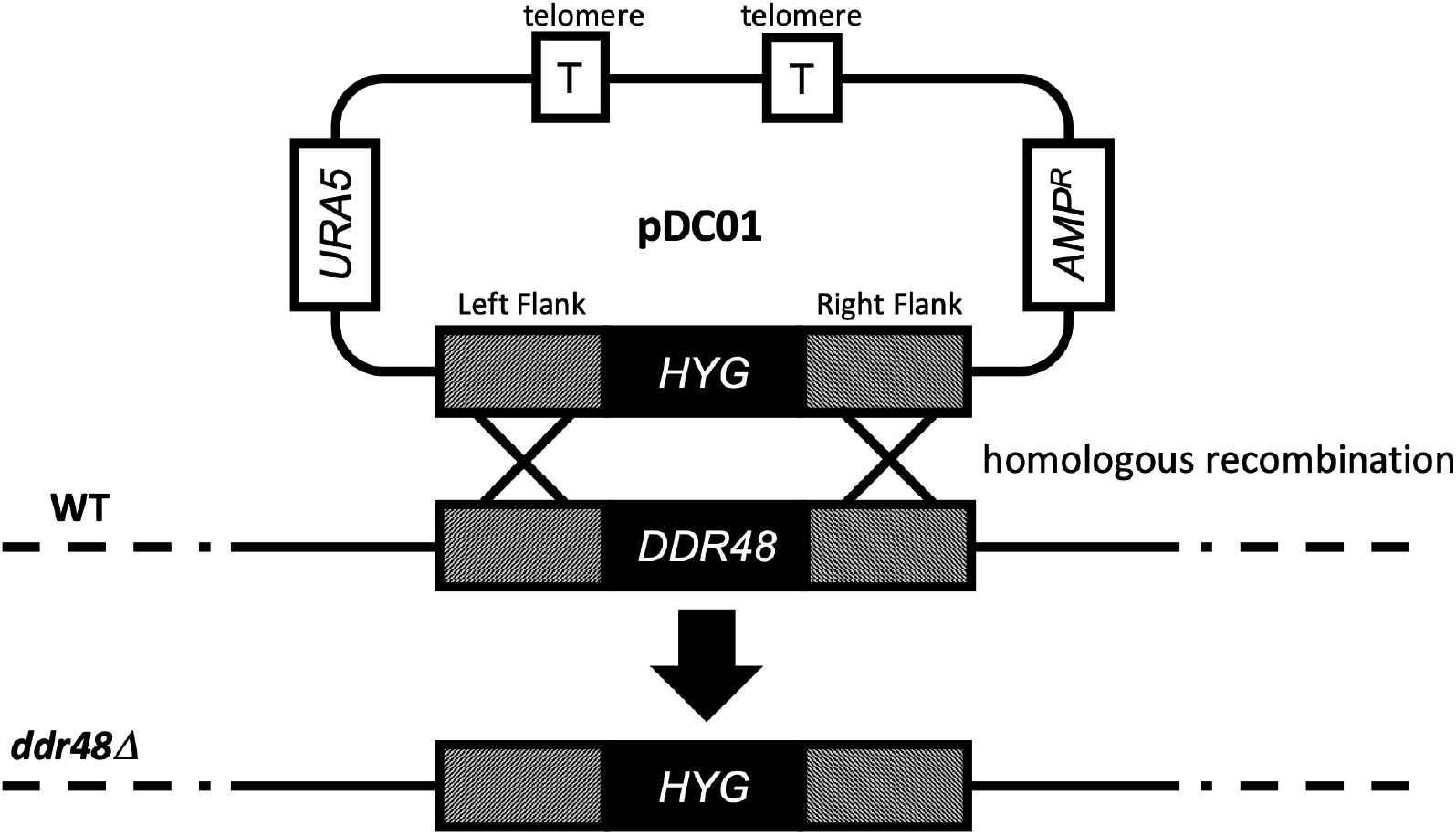
Schematic of *DDR48* Allelic Replacement. Depiction of homologous recombination of the hygromycin resistance locus jointed between *DDR48* gDNA and the native *DDR48* genomic sequence.

**Figure S5:**
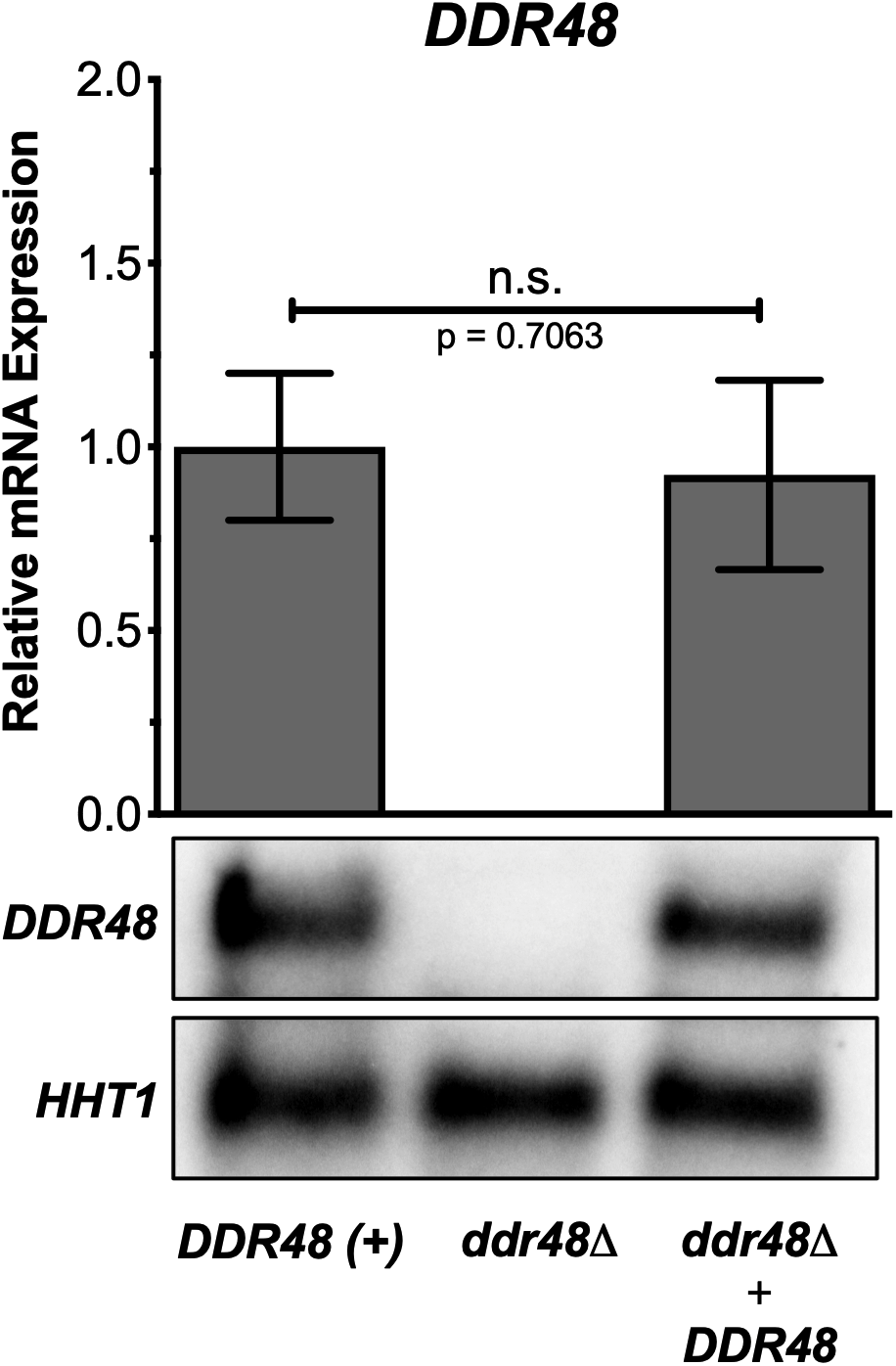
Quantitative and qualitative confirmation of *DDR48* deletion. qRT-PCRs were performed on wildtype, *ddr48*Δ allelic replacement mutant, and *ddr48 + DDR48* complement strains. *Histone H3* was used as a normalizer gene using the ΔΔ_ct_ method to normalize to wild-type *DDR48 (+)* expression. Northern blot analysis were performed on wildtype, *ddr48*Δ allelic replacement mutant, and *ddr48* + *DDR48* complement strains. All data generated were performed with at least two biological replicates. Data from replicates are graphed as mean ± standard deviation

**Figure S6:**
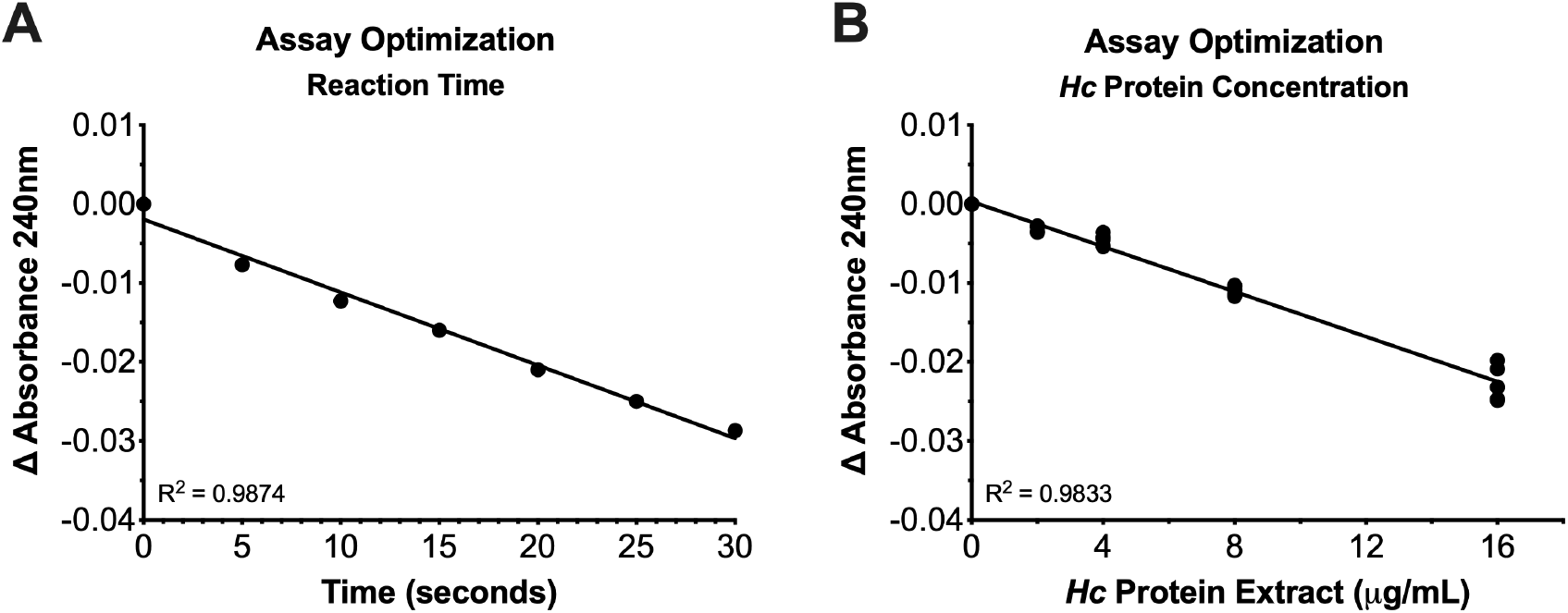
Standard curve optimization of peroxide destruction assay. Reaction time **(A)** and protein concentration **(B)** standard curves used to determine optimized reaction conditions for the hydrogen peroxide destruction assay.

**Table S1:** *Histoplasma* strains used in this study. ^a^all strains were constructed from G186A background strain ^b^“smooth” colony variant and uracil auxotroph of G186A (CIT)

| Strain<sup>a</sup> | Genotype | Other Designation |
| --- | --- | --- |
| G186A | Wildtype (ATCC# 26029) |  |
| WU27<sup>b</sup> | *ura5Δ* | WT; *DDR48* (+) |
| USM10 | *ura5Δ ddr48Δ::hph* | *ddr48Δ* |
| USM13 | *ura5Δ ddr48Δ::hph / pLE4 [URA5, DDR48]* | *ddr48Δ + DDR48* |

**Table S2:** *BLAST* analysis of fungal *DDR48* protein. Fungal BLAST table of the *H. capsulatum DDR48* amino acid sequence from the *Saccharomyces* Genome Database (SGD) Fungal BLAST Suite.

| Species | Max Score | Total Score | Query Coverage | E-value | Percent Identity |
| --- | --- | --- | --- | --- | --- |
| *Blastomyces dermatitidis* | 282 | 282 | 93% | 1.00E-88 | 60.31% |
| *Candida albicans* | 109 | 181 | 79% | 2.00E-24 | 52.76% |
| *Candida glabrata* | 98 | 482 | 80% | 4.00E-19 | 46.07% |
| *Coccidioides immitis* | 101 | 101 | 98% | 2.00E-20 | 35.73% |
| *Emmonsia crescens* | 259 | 372 | 99% | 3.00E-80 | 64.63% |
| *Microsporium canis* | 125 | 125 | 96% | 9.00E-30 | 42.74% |
| *Paracoccidioides brasiliensis* | 127 | 127 | 92% | 9.00E-31 | 48.51% |
| *Penicillium brasilianum* | 115 | 188 | 91% | 2.00E-26 | 49.33% |
| *Saccharomyces cerevisiae* | 106 | 584 | 81% | 7.00E-22 | 48.60% |

**Table S3:**
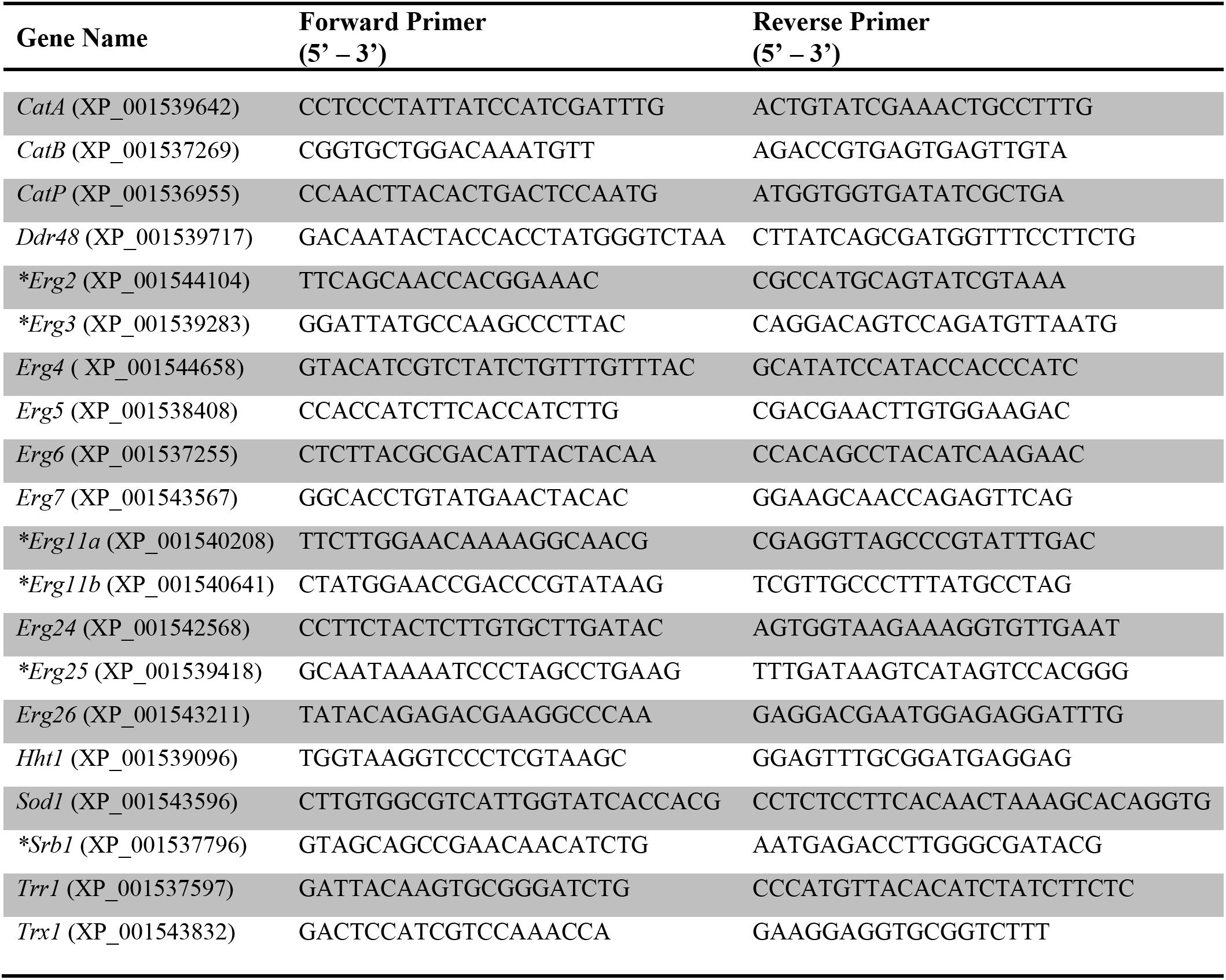
qRT-PCR primer sequences. * = primer sequence adapted from DuBois JC, Smulian AG (2016) Sterol Regulatory Element Binding Protein (Srb1) Is Required for Hypoxic Adaptation and Virulence in the Dimorphic Fungus *Histoplasma capsulatum*. PLOS ONE 11(10): e0163849.

## Materials and Methods

### Strains and culture conditions

All *H. capsulatum* strains used in this study were derived from the wildtype strain G186AR (ATCC 26029), which are listed in **Table S1** in the supplemental material. *Histoplasma* yeasts were cultured in *Histoplasma* macrophage medium (HMM), a modified tissue culture medium optimized to mimic *Histoplasma* growth *in vivo* (65). All cultures were supplemented with ampicillin and streptomycin to decrease bacteria contamination over longer growth periods. *Histoplasma* strains auxotrophic for uracil were supplemented with uracil (100 ug/ml). *Histoplasma* yeasts were grown at 37°C with shaking at 200 rpm until mid/late log phase before performing experiments. Growth of *Histoplasma* yeasts was determined by culture turbidity at 600 nm. For measurement of growth kinetics, *Histoplasma* yeasts were grown to mid/late log phase then diluted with pre-warmed, pre-aerated HMM to a final turbidity at 600 nm = 0.1 and incubated at 37°C with shaking at 200 rpm for a duration of 7 days. Turbidity measurements were taken every 24 hours to assess growth.

### Generation of the *ddr48* mutant and *DDR48*-complemented strains

The *DDR48* deletion strain was previously created in our laboratory using allelic replacement as described previously (66). A schematic of the allelic replacement design can be found in **SI4**. For complementation of *DDR48*, the *DDR48* gene, and 1,000 bp upstream to encompass the *DDR48* promoter region, of start was PCR amplified from parent strain WU27 genomic DNA and cloned into pRPU1, a telomeric expression vector optimized for *Histoplasma* genetics. The *ddr48Δ* mutant was then transformed with the *DDR48*-expressing telomeric vector by electroporation and URA^+^ transformants were screened for *DDR48* gene expression.

### DNA and RNA Extractions and Blotting

DNA and RNA were extracted from mid/late log phase *Histoplasma* yeasts as previously described (67). Briefly, cells were harvested by centrifugation and re-suspended in RNA extraction buffer (0.1M sodium acetate, pH 5.0, 0.2M NaCl, 0.2% w/v sodium dodecyl sulfate) or DNA extraction buffer (0.1M TRIS, pH 8.0, 0.1M EDTA, 0.25M NaCl) along with 0.5 mm acid washed glass beads and phenol:chloroform (5:1). The nucleic acids were then liberated by bead-beating until roughly 80% of cells were lysed as determined by microscopy. The nucleic acids were then precipitated from the aqueous fraction by the addition of 2 volumes of pure ethanol and stored at −20°C until use. Northern blotting was performed as previously described (35).

### Quantitative Real-Time PCR

Primer pairs for each gene target used in this study were designed to yield an approximate 200 bp product, which can be found in **Supplemental Table 3**. RNA was extracted per the protocol listed above. RNA was subjected to DNAse treatment prior to cDNA synthesis using TURBO DNA-free kit (Invitrogen). A total of 500 ng of RNA was then reverse-transcribed with the Maxima First Strand cDNA Synthesis Kit for qRT-PCR, with dsDNase (Thermo Fisher) per the manufacture’s protocol. The cDNA was quantified via A260/A280 using a NanoDrop (Fisher Scientific) spectrophotometer and diluted to a final concentration of 500 ng/μl. The diluted cDNA was then stored at 4°C until needed.

Quantitative, real-time PCR (qRT-PCR) was performed on triplicate samples containing 500 ng of reverse-transcribed RNA using the Maxima SYBR Green/ROX qPCR Master Mix (2X) (Thermo Fisher). Each reaction contained 0.2 μM forward primer, 0.2 μM reverse primer, 12.5 μl 2X SYBR Green/ROX Master Mix, 500 ng of cDNA, and nuclease-free water to a total volume of 25 μl. To reduce pipetting errors, a master mix was assembled for each gene-specific primer set containing all reagents except cDNA and dispensed in to each tube. 1 μl of 500 ng/μl cDNA was then individually added to each tube before starting the reaction. The qRT-PCR reactions were performed using a CFX96 Touch™ Real-Time PCR Detection System using the following conditions: 95°C for 10 minutes followed by 40 cycles of 95°C for 15 seconds, 60°C for 30 seconds, 72°C for 30 seconds. Integrity of each run was measured by melt-curve analysis. Relative expression was determined using the *ΔΔ*_*Ct*_ method after normalizing to levels of the constitutively expressed house-keeping gene *Histone H3 (HHT1)* transcript.

### Susceptibility to superoxides and peroxides

Yeast phase *Histoplasma* cultures were grown to mid-log phase. The cultures were then diluted with pre-warmed, pre-aerated, HMM to an OD_600_ = 0.1. To generate the superoxide anions, a 20 mM stock of paraquat dichloride was used to bring 50-ml aliquots of the diluted *Hc* cultures, in triplicate, to a final concentration of 0.5 μM, 0.75 μM, and 1.0 μM, respectively. Immediately before performing the experiment, a 33% v/v hydrogen peroxide solution (Sigma) was diluted with 1X PBS to make a fresh 50 mM hydrogen peroxide stock solution. The 50 mM stock was then used to bring 50-ml aliquots of diluted *Hc* cultures, in triplicate, to a final concentration of 2.5 mM, 5.0 mM, and 7.5 mM, hydrogen peroxide, respectively. Controls were also prepared by taking triplicate 50-ml aliquots of diluted *Hc* cultures with no additions. The cultures were then incubated at 37°C in a humidified chamber with shaking at 200 rpm. Every 24-hours, 1-ml of each culture was removed and added to a plastic 1.5 ml polystyrene cuvette and sealed using a polyethylene cuvette cap. The cuvettes were slowly inverted three times to homogenously mix each sample right before being placed in to the cuvette reader. The absorbance at 600 nm was measured three times for each sample and recorded. Measurements were taken every 24 hours for a duration of 5 days (120 hours). Absorbance measurements at OD_600_ were taken every 24 hours.

### Catalase and superoxide dismutase (SOD) assays

The superoxide dismutase (SOD) assay was performed per the manufacturer’s protocol (Cayman Chemical). The hydrogen peroxide destruction assay was performed using a modified protocol previously described (16, 95) to extrapolate relative catalase activity. In brief, 33% hydrogen peroxide (Sigma) was diluted to a final concentration of 10 mM PBS (137 mM NaCl, 10 mM Na_2_HPO_4_, 2.7 mM KCl, and 1.8 mM KH_2_HPO_4_; pH 7.4), added to a clean 100-ml glass bottle, and placed on ice immediately before performing the experiment.

Standard curves were generated to determine optimal protein concentration and incubation time for the assay **SI6**. Eight micrograms of total protein lysate from each *Hc* strain was diluted to a final volume of 200 μl using PBS and allowed to equilibrate to 4°C on ice. The reaction was assembled right before use by adding 800 μl of the 10 mM hydrogen peroxide solution to the 200 μl *Hc* protein extract, mixing briefly so not to create bubbles, and placed into a 1 ml quartz cuvette (Sigma). The quartz cuvette was then immediately placed into the cuvette holder of a microplate reader (Molecular Devices) where the absorbance at 240 nm and 600 nm was recorded every 20 seconds for a total of 5 minutes. The above methods were subsequently repeated for each individual *Hc* protein extract being examined right before reading the absorbance to ensure the linear phase of catalase activity occurred in the measured time frame of the assay.

### Susceptibility to antifungal agents

Susceptibility of *Hc* strains to the antifungal drugs amphotericin b and ketoconazole was determined by using a microplate-based growth assay previously described (68). Briefly, *Histoplasma* yeasts were diluted in a total volume of 10-mls pre-warmed 2X HMM and 50 μl aliquots were added to each well of a 96-well, flat-bottomed microplate. Next, 50 μl of water only or water supplemented with amphotericin b or ketoconazole, respectively, at twice the desired concentration (two-fold dilutions from 32 μg/ml to 0.03 μg/ml final concentrations) was added to each well for a total volume equaling 100 μl. Each well was mixed gently using a multichannel micropipette by pipetting up-and-down for 10 seconds carefully, as to not create bubbles. The lid was placed on the microplate and sealed with Blenderm™ (3M) breathable tape. The plates were then incubated for at 37°C in a humidified chamber with twice daily aeration by incubating plates on a bench-top rocker for 30 minute intervals every 12 hours. The absorbance of each well at 600 nm was measured using a microplate reader on the fourth day of the experiment. The measurements were then entered in to GraphPad Prism software where IC_50_ was calculated by nonlinear regression.

### Macrophage infections

RAW 264.7 macrophages (ATCC TIB-71), *Mus musculus*, were thawed from a frozen stock, centrifuged at 500 x g for 5 minutes in a swing-bucket centrifuge, and re-suspended in an equal volume of fresh, pre-warmed 1X DMEM supplemented with 10% v/v FBS, 50 μg/ml ampicillin, and 100 μg/ml streptomycin. The aliquot was then added to a T-75 cell culture flask containing 20-ml of 1X DMEM, with supplements mentioned above, and incubated at 37°C in a humidified incubator with 5% CO_2_ and 95% room air atmosphere for 24 hours. Once the culture reached ~80% confluence, the macrophages were dissociated with 0.25% trypsin-EDTA (Gibco), enumerated by microscopy with a hemacytometer, and 5 × 10^4^ macrophages were seeded into each well of a 24-well tissue culture plate in a total volume of 1-ml. For assays that required macrophage activation, the macrophages were allowed to adhere to the culture plate for 20 minutes before 100 units (U) of murine recombinant interferon-gamma (IFNγ) (Invitrogen) was added to each well to stimulate the macrophages. The seeded 24-well tissue culture plates were then incubated at 37°C in a humidified incubator with 5% CO_2_ and 95% room air atmosphere for 24 hours to allow the cells to grow to confluence. Once the macrophages reached ~80% confluence, the wells were washed with 1X DPBS three times. For co-infection, *Histoplasma* yeasts were enumerated via a hemacytometer and diluted to 2 × 10^5^ cells/well, 2 × 10^6^ cells/well, or 1 × 10^7^ cells/well, which corresponds to a multiplicity of infection (MOI) (macrophage-to-yeast) ratio of 1:1, 1:10, or 1:50, respectively, in HMM-M (HMM supplemented with 10% FBS, 584 mg/L L-glutamine, and 3.7 g/L sodium bicarbonate). The macrophage-yeast co-cultures were then incubated at 37°C in a humidified incubator with 5% CO_2_ and 95% room air atmosphere for 2 hours to facilitate phagocytosis. Each well was then subsequently washed with 1X DPBS three times to remove any un-phagocytized *Hc* yeasts. The wells were then re-suspended in an equal volume of fresh, pre-warmed HMM-M and incubated at 37°C in a humidified incubator with 5% CO_2_ and 95% room air atmosphere. At 24-hours post-infection, the culture medium was removed from each well, washed with DPBS three times, and an equal volume of water added to facilitate macrophage lysing. The macrophages were mechanically lysed by scraping the bottom of each well with a sterile micropipette tip. Dilutions of each lysate were plated on to triplicate HMM plates and incubated at 37°C until visible colonies appeared. Yeast survival was determined by viable colony forming units (CFUs) as a percentage survival of un-activated, recovered colonies.

### Phagocytosis assay

Phagocytosis assays were performed using a modified protocol from Cordero et al. (69). RAW 264.7 macrophages were plated onto 6-well tissue culture plates containing an 18 mm diameter glass coverslip coated in poly-L-lysine at 1.5 × 10^5^ cells per well in a total volume of 3-ml of 1X DMEM. The macrophages were incubated for 24-hours at 37°C in a humidified incubator with 5% CO_2_ and 95% room air atmosphere to allowed adherence to the coated glass coverslip. *Histoplasma* yeast cultures were grown in HMM at 37°C to mid-log growth phase before 10-ml aliquots were removed and labelled with 40 μg/ml of NHS Rhodamine (Thermo-Fisher) for 30 minutes at 25°C. The cells were then washed with an equal volume of 1X DPBS three times before re-suspending in an equal volume of pre-warmed HMM and enumerated by a hemacytometer. For co-infection, the media was removed from each well of macrophages, washed with 1X DPBS three times, and 7.5 × 10^5^ *Hc* cells (MOI = 1:5) were added to each well in a total volume of 3-ml of pre-warmed HMM-M. The plates were incubated for two hours at 37°C in a humidified incubator with 5% CO_2_ and 95% room air atmosphere to facilitate phagocytosis of the labelled *Hc* yeasts. After the incubation, the plates were washed with 1X DPBS three times, fixed for 15 minutes at 25°C with 3-mls of 4% formaldehyde solution in PBS, and washed with 1X DPBS three times. Each coverslip was then removed from their well, placed onto a clean glass microscope slide containing 4 μl of hard set mounting medium (Vecta-Shield), and allowed to set for 10 minutes before sealing with nail polish. The slides were allowed to dry in the dark overnight at 25°C before being stored long term in a slide box at 4°C until microscopic analysis. Images were obtained using a Zeiss 510 Meta confocal microscope using DIC and the 650 nm laser using immersion oil at 60X magnification (Supplemental Figures). At least 8 different images were obtained, each of a different field of view. The number of macrophages and the number of *Hc* yeasts within each were recorded for each field of view. At least 200 macrophages were counted for each *Hc* strain being analyzed. The percentage of phagocytosis was determined by calculating the ratio of macrophages containing *Hc* yeasts divided by the total number of macrophages enumerated. The phagocytic index was determined by calculating the average number of *Hc* yeasts inside of the recorded macrophages.

### Statistical Analysis

All data generated were performed on three technical replicates and at least two biological replicates and represented as mean ± SEM. Data were analyzed by students t test, one-way analysis of variance (ANOVA), or two-way ANOVA followed by Tukey’s multiple comparisons test using GraphPad Prism v8. All p-values are included in each figure.

## Acknowledgements

We thank Davida Crossley for construction of the *ddr48Δ Hc* strain previously in the Shearer laboratory. We are grateful to Jonathan Lindner and the MS INBRE Imaging Facility for equipment use and helpful discussions regarding data collection. We thank Erin Walker and Paige Brady for assistance in screening *DDR48* complement strains and running qRT-PCR plates, respectively. We also thank William Goldman (UC Chapel Hill) for providing the WU27 *Hc* strain and George S. Deepe, Jr. for technical editing of the manuscript.

